# A Pairwise Distance Distribution Correction (DDC) algorithm to eliminate blinking-caused artifacts in super-resolution microscopy

**DOI:** 10.1101/768051

**Authors:** Christopher H. Bohrer, Xinxing Yang, Xiaoli Weng, Brian Tenner, Shreyasi Thakur, Ryan McQuillen, Brian Ross, Matthew Wooten, Xin Chen, Melike Lakadamyali, Jin Zhang, Elijah Roberts, Jie Xiao

## Abstract

In single-molecule localization based super-resolution microscopy (SMLM), a fluorophore stochastically switches between fluorescent- and dark-states, leading to intermittent emission of fluorescence, a phenomenon known as blinking. Intermittent emissions create multiple localizations belonging to the same molecule, resulting in blinking-artifacts within SMLM images. These artifacts are often interpreted as true biological assemblies, confounding quantitative analyses and interpretations. Multiple methods have been developed to eliminate these artifacts, but they either require additional experiments, arbitrary thresholds, or specific photo-kinetic models. Here we present a method, termed Distance Distribution Correction (DDC), to eliminate blinking-caused repeat localizations without any additional calibrations. The approach relies on the finding that the true pairwise distance distribution of different fluorophores in an SMLM image can be naturally obtained from the imaging sequence by using distances between localizations separated by a time much longer than the average fluorescence survival time. We show that using the true pairwise distribution we can define and then maximize the likelihood of obtaining a particular set of localizations void of blinking-artifacts, generating an accurate reconstruction of the underlying cellular structure. Using both simulated and experimental data, we show that DDC surpasses all previous existing blinking-artifact correction methodologies, resulting in drastic improvements in obtaining the closest estimate of the true spatial organization and number of fluorescent emitters in a wide range of applications. The simplicity and robustness of DDC will allow it to become the field standard in SMLM imaging, enabling the most accurate reconstruction and quantification of SMLM images to date.

## Introduction

In recent years the development of superresolution fluorescence microscopy has enabled the probing of macromolecular assemblies in cells with nanometer resolutions. Amongst different superresolution imaging techniques, single-molecule localization superresolution microscopy (SMLM) has gained wide popularity due to its relatively simple implementation, which is based on post-imaging analysis of single-molecule detection.

SMLM reconstructs a superresolution image by stochastic photo-activation and subsequent post-imaging localization of single fluorophores (1–3). One major advantage of SMLM is that due to its single-molecule detection nature, one can determine the number of molecules in a macromolecular assembly quantitatively, allowing the investigation of both the molecular composition and spatial arrangement at a level unmatched by other ensemble imaging-based superresolution imaging techniques. In the past few years SMLM has led to novel discoveries and quantitative characterizations of numerous biological assemblies (4, 5) such as those composed of RNA polymerase (6–8), membrane proteins (9), bacterial divisome proteins (10–13), synaptic proteins (14, 15), the cytoskeleton (16), DNA binding proteins (17, 18), chromosomal DNA (19), viral proteins (20), and more.

One critical aspect in realizing the full quantitative potential of SMLM relies on the careful handling of the blinking behavior of fluorophores. A photo-switchable fluorophore can switch multiple times between activated and dark states before it is permanently photobleached, leading to “repeat localizations” from the same molecule. These repeat localizations are often misidentified as multiple molecules, adding additional levels of error to the superresolution images. For example, blinking-artifacts often lead to the formation of false nanoclusters and errors in quantifying numbers of molecules and the stoichiometry of complexes (Fig. 1A) (21–25).

**Figure 1:**
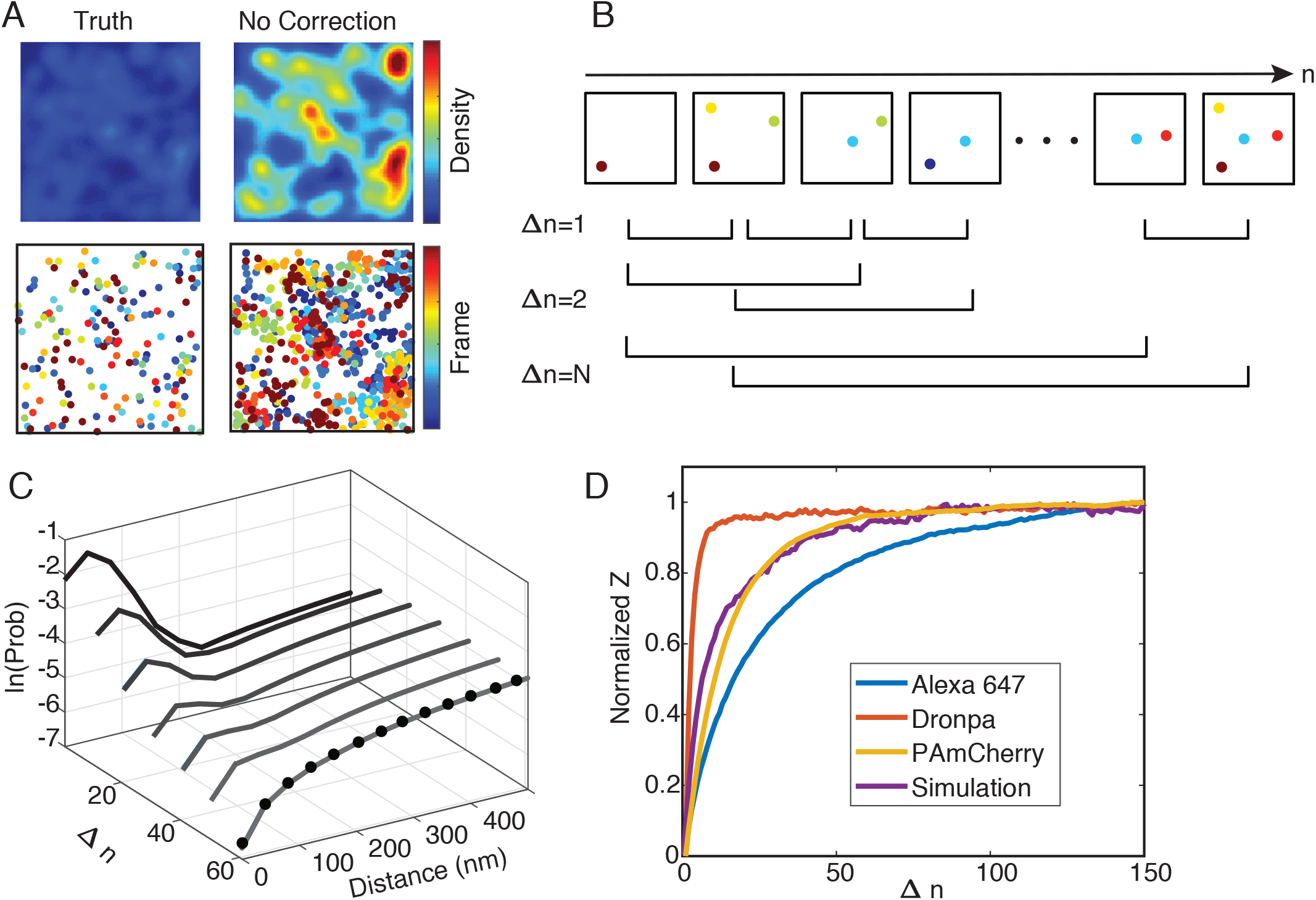
A. Simulated SMLM superresolution images (top panel) of randomly distributed molecules without repeats (Truth) and with repeats (No correction). The corresponding scatter plots (colored through time) are displayed in the bottom panel. B. Schematics of how the pairwise distance distributions at different frame differences (Δ*n*) were calculated. C. Pairwise distance distributions at different Δ*n* (black to gray curves) converge to the true pairwise distribution (black dots) when Δ*n* is large. D. Normalized Z values measured for three commonly used fluorophores and a simulated fluorophore as that used in A. All Z values reach plateaus at large Δ*n*, indicating that at large Δ*n*, the pairwise distance distributions converge to a steady state. The normalized Z value was calculated by taking the difference between the cumulative pairwise distance distribution at a Δ*n* and that at Δ*n* = 1: (*Z*(Δ*n*) = Σ|*cdf*(*P_d_*(Δ*r*|Δ*n*)) − *cdf*(*P_d_*(Δ*r*|Δ*n*= 1))|).

Multiple groups have developed different methods to correct for blinking-caused artifacts in SMLM. These methods can be coarsely divided into two categories depending on whether a method provides a corrected image void of repeat localizations or a statistical analysis summarizing the properties of the image at the ensemble level. Methods in the first category commonly use a variety of threshold values both in time and space to group localizations that likely come from the same molecule (1, 2, 21, 23, 25, 26). The advantage of using thresholds is that it results in a corrected image, allowing one to observe the spatial distribution of fluorophores in cells and apply other quantitative analyses as needed. The disadvantage is that a constant threshold value is often insufficient in capturing the stochastic nature of fluorophore blinking and heterogeneous molecular assemblies. Furthermore, calibration experiments and/or a priori knowledge of the fluorophore’s photochemical properties are often needed to determine the appropriate threshold values (21, 23, 25, 27, 28). Statistical analyses such as maximum likelihood or Bayesian approaches have been developed to take into account the stochastic behavior of blinking but have yet to produce corrected superresolution images void of repeat localizations (29–31). Additionally, many of these approaches are dependent on specific photokinetic models for the fluorophore, which can be complex and difficult to determine (27, 28, 32–35).

The second category of methods use statistical methods to characterize mean properties of the organization of molecules at the ensemble level in raw, uncorrected SMLM images. Pair- or auto-correlation-based analyses (PCA) have been used extensively in the field (24, 36). The long tail of the correlation function can often be fit to a specific model to extract quantitative parameters. This class of methods is prone to model-specific errors, especially if the underlying structures of the molecular assemblies are heterogeneous and vary throughout the image (37). A recently developed method analyzes the clustering of a protein with experimentally varied labeling densities, which was robust in determining whether membrane proteins form nanoclusters and was insensitive to many imaging artifacts (22). A post-imaging computational analysis capitalizing on the same principle has also been developed (38). Although these methods are powerful in determining whether a protein of interest forms clusters or not, they provide a quantification at the ensemble level but not a corrected image, which limits their use in analyzing heterogeneously distributed molecular assemblies and their spatial arrangement in cells.

Here, we present an algorithm, termed Distance Distribution Correction (DDC), to enable robust reconstruction and quantification of SMLM superresolution images free of blinking-caused artifacts without the need of setting empirical thresholds or performing calibration experiments. We first validate our approach using a diverse set of simulated and experimental data and compare DDC to other existing methods. In each situation DDC outperformed the existing methods in obtaining the closest representation of the “true” image and in determining the accurate number of fluorophores. We then applied DDC to experimentally collected SMLM images of membrane scaffolding proteins (46–48), dynein oligomers (39) and isolated sister chromatin fibers (40). Under all the conditions tested, DDC provided SMLM superresolution images devoid of repeat localizations caused by fluorophore blinking, allowing identification of membrane protein cluster properties, characterizations of dynein in different assembly states, and quantification of DNA content between sister chromatin fibers. These results demonstrate the broad application of DDC for SMLM imaging. Finally, we discuss critical considerations of how to apply DDC to experiments successfully.

## Results

### Principle of DDC

DDC is based on the principle that the pairwise distance (Δ*r*) distribution, *P_d_*(Δ*r*|Δ*n*), of the localizations separated by a frame difference (Δ*n*) much larger than the average number of frames a molecule’s fluorescence lasts (*N*) approximates the true pairwise distance distribution *P_T_*(Δ*r*). Note that *N* does not need to be precisely determined as long as it is in the regime where *P_d_*(Δ*r*|Δ*n*) approaches a steady state, as we show below. One intuitive way to understand this principle is that, if one collects an imaging stream that is long enough so that all the localizations in the first and last frames of the stream come from distinct sets of fluorophores, the pairwise distance distribution between the localizations of the two frames will then be devoid of repeat localizations and will reflect the true pairwise distance distribution (*P_T_*(Δ*r*)). A mathematical justification of this principle is provided in the supplemental material with an in-depth discussion and illustration (Fig. S1).

To demonstrate the principle of DDC, we used simulated SMLM images of randomly distributed fluorophores that followed the photokinetic model shown in Fig. S2A. One representative superresolution image and the corresponding scatter plot, colored through time, with and without repeat localizations are shown in Fig. 1A. Apparent clustering was observed in images when repeat localizations were not corrected. Using the uncorrected images, we computed the pairwise distance distributions at all frame differences Δ*n* (Fig. 1B). As shown in Fig. 1C and Fig. S3, at small Δ*n* there are large peaks at short distances, indicating that there were repeat localizations from the same fluorophores closely spaced in time and space. When Δ*n* is large, the pairwise distance distributions approach a steady state converging upon the true pairwise distance distribution (Fig. 1C, dotted curve). This behavior supports the principle that when Δ*n* is sufficiently large the pairwise distance distribution represents the true pairwise distance distribution. Using simulations, we also show that the pairwise distance distributions converge upon the true distributions at large Δ*n* irrespective of the underlying photokinetics or molecular spatial distributions (Fig. S3, Supporting Material).

Next, we used experimentally obtained SMLM images of three molecular assemblies labeled with different fluorophores in *E. coli* cells, the bacterial transcription elongation factor NusA fused with the reversibly switching green fluorescent protein Dronpa (41), *E. coli* RNA Polymerase fused with the photoactivatable red fluorescent protein PAmCherry (42), and precursor ribosomal RNAs (pre-rRNA) labeled with organic fluorophore Alexa647-conjugated DNA probes (43) (Fig. S4, Supporting Material). We determined the pairwise distance distribution for each fluorophore and calculated the normalized, summed differences of the cumulative distributions for each Δ*n*, relative to that of Δ*n* = 1, (*Z*(Δ*n*) = Σ|*cdf*(*P_d_*(Δ*r*|Δ*n*)) – *cdf*(*P_d_*(Δ*r*|Δ*n* = 1))|). As shown in Fig. 1D, in all cases the corresponding normalized *Z* values reach plateaus at large Δ*n* despite different photokinetics and spatial distributions. The rate at which each fluorophore reaches the plateau for the normalized *Z* reflects the photokinetics of the fluorophore — the longer a fluorophore blinks (such as Alexa647 compared to Dronpa), the longer the time until *Z* plateaus. These experimental results further confirm the principle of DDC that the pairwise distance distributions converge upon a steady state distribution as Δ*n* increases.

It is important to note that the determination of *P_T_*(Δ*r*) is not dependent upon a particular photokinetic model of the fluorophore nor does it require experimental characterizations of the fluorophore. *P_T_*(Δ*r*) can be determined solely from the SMLM image stream as long as it is long enough so that a steady state of *P_d_*(Δ*r*|Δ*n*) can be reached (Fig. 1C, Fig. S3).

Once determined, *P_T_*(Δ*r*) can then be used to calculate the likelihood to have a particular subset of true localizations (Fig. S5-S9, Supporting Material) using the following equation:

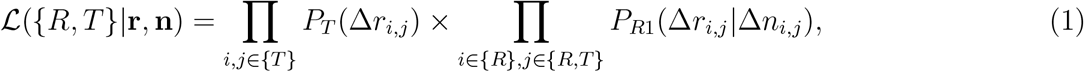

where {*R,T*}are sets that contain the indices of the localizations that are considered repeats {*R*} and the true localizations {*T*}given the coordinates **r** and associated frame numbers **n** obtained from experiment. The first term on the right of the equation is the probability of observing all distances Δ*r* between every pair of true localizations (*i* & *j* ∈ {*T*}). Here the probability distribution *P_T_*(Δ*r_i,j_*) is the true pairwise distance distribution. The second term is the probability of observing all distances between pairs of localizations with at least one being a repeat (*i* ∈ {R} and *j* ∈ {*R,T*}). Here, the probability distribution *P*_*R*1_(Δ*r_i,j_*|Δ*n_i,j_*) gives the probability of observing a distance between a pair of localizations with a frame difference Δ*n_i,j_* if at least one of the localizations is a repeat. This probability distribution can be easily determined once *P_T_*(Δ*r*) is known (Supporting Material). Here, maximizing the likelihood with respect to {*R,T*} results in a subset of true localizations where the pairwise distance distributions *P_d_*(Δ*r*|Δ*n*) are equal to *P_T_*(Δ*r*) (Fig. S6). DDC maximizes the likelihood with respect to the two sets ({R, T}) using a Markov Chain Monte Carlo (MCMC) (44, 45) to reconstruct the corrected image (Fig. S8 and S9, Supporting Material).

To validate Equation 1, we performed six simulations of distinct spatial distributions with various fluorophore photo-kinetic models. We found that only when greater than 97% of the final localizations were true localizations did the likelihood reach its maximum (Fig. S7).

### DDC outperforms existing methods in both image reconstruction and quantifications

To compare the performance of DDC with commonly used blinking-artifact eliminating methods, we simulated five systems, random distribution (no clustering), small clusters, dense clusters, parallel filamentous structures with low labeling density, and intersecting filamentous structures with high labeling density (Fig. 2, Supporting Material). In these simulations the fluorophore had two dark states and followed the photokinetic model shown in Fig. S2A. The raw images without any repeat localizations for each simulation are shown in Fig. 2A. We applied DDC, three published thresholding methods (T1 to T3 (21, 23, 25))(Supporting Material, Fig. S10 and S11) and a customized thresholding method (T4, Supporting Material) to all the images.

**Figure 2:**
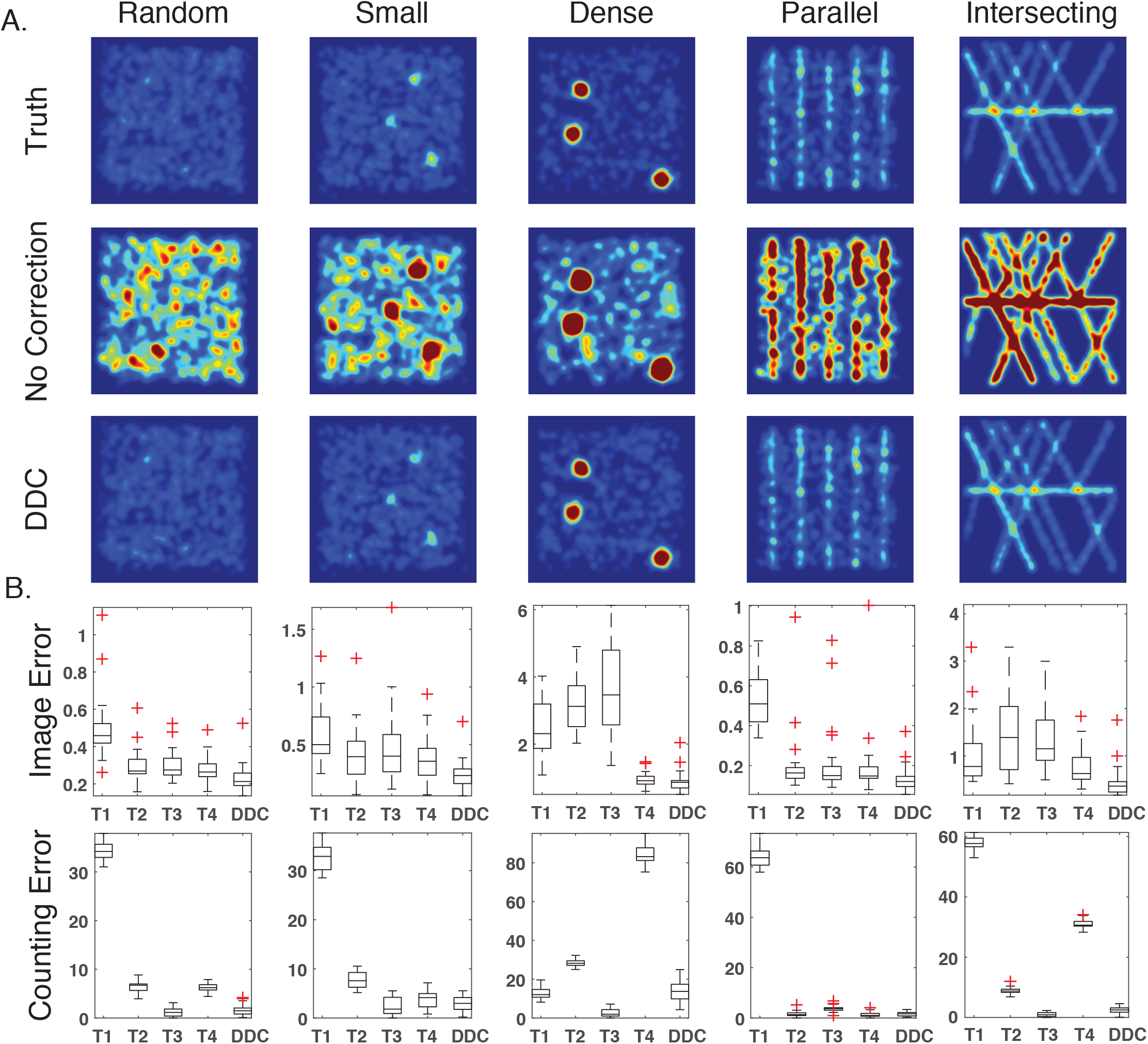
Comparison of four different thresholding methods with DDC on five spatial distributions (randomly distributed, small clusters, dense clusters and parallel filaments and intersecting filaments). A. True, uncorrected and DDC-corrected images for each spatial distribution. B. Image Error and Counting Error calculated from T1 to T4 and DDC for each spatial distribution. The whiskers extend to the most extreme data points not considered outliers, and the red pluses are the outliers (greater than 2.7 std).

Method T1 links together localizations using a time threshold that is determined by an empirical estimation of the photokinetics of the fluorophore (21) (Fig. S10, Supporting Material). Method T2 uses experimentally quantified photo-kinetics of the fluorophore to set extreme thresholds so that the possibility of overcounting is extremely low (25). Method T3 uses the experimentally determined number of repeats per fluorophore to choose thresholds that result in the correct number of localizations within each image (23)(Fig. S11, Supporting Material). T2 and T3, but not T1, require additional experiments to characterize fluorophore photo properties. Method T4 is a customized, ideal thresholding method that scans all possible thresholds and uses the threshold that results in the least Image Error for each system (Supporting Material). T4 cannot be applied in real experiments since the true, repeat localization-free image is unknown — we included it here to illustrate the best scenario of what a thresholding method could achieve.

To quantitatively compare the ability of these methods in producing a repeat localization-corrected image we calculated two metrics, the Image Error and Counting Error (Fig. 2B, Supporting Material). The Image Error was calculated by first summing the squared difference of each pixel’s normalized intensity between the corrected and the true images, and then dividing this squared difference by the error between the uncorrected and the true images (Supporting Material). The Image Error quantifies the amount of error in determining the distribution of localizations without being penalized for the error in the number of localizations. The Counting Error was calculated as the difference between the true number of fluorophores and that determined from the corrected image divided by the actual number of fluorophores (Supporting Material).

As shown in Fig. 2B, DDC outperforms all four methods by having the lowest Image Errors and lowest (or close-to-lowest) Counting Errors. Interestingly, even with the best possible thresholds (T4), DDC still outperforms T4 in determining the correct spatial distribution and numbers of localizations. This result suggests that thresholds cannot adequately account for the stochastic nature of blinking. Similar results are shown in Fig. S12 for a fluorophore with one dark state (Fig. S2B). When counting the number of localizations is the main concern, T3 performs equally or slightly better than DDC because T3 was applied with an experimental calibration that provides the average number of blinks per fluorophore (Fig. 2, Supporting Material). Nonetheless, DDC outperforms T3 by having lower Image Errors across all five simulation systems of different spatial organization patterns. In particular, for the dense cluster and the intersecting filament systems, two scenarios commonly encountered in biology for the spatial organizations of membrane and cytoskeletal proteins, the average Image Errors of T3 are more than four times that of DDC (Fig. 2B). The significant advantage of DDC over other methods for these two systems particularly highlights the unique superiority of DDC in handling heterogeneously distributed proteins with uneven densities due to clustering or overlapping structures. In conclusion, these results demonstrate that DDC can be used to obtain the correct number of true localizations and at the same time produce the most accurate SMLM images.

### DDC identifies differential clustering properties of membrane microdomain proteins AKAP79 and AKAP150

Membrane microdomains formed by membrane proteins have been commonly observed in super-resolution imaging studies and have raised significant interest in their molecular compositions and associated biological functions (9). However, concerns remain as of whether the characterizations of these microdomain protein clusters were impacted by blinking-caused artifacts (22). Here we used DDC to investigate a membrane scaffolding protein, A-Kinase Anchoring Protein (AKAP), which plays an important role in the formation of membrane microdomains (46–48). The two orthologs AKAP79 (human) and AKAP150 (rodent) were previously shown to form dense membrane clusters, which are likely important for regulating anchored kinase signaling.

We performed SMLM imaging on AKAP150 in murine pancreatic beta cells using an anti-AKAP150 antibody and analyzed the resulting SMLM data using DDC (Supporting Material). For AKAP79, we applied DDC to previously acquired SMLM data from HeLa cells (46). For comparison, we also applied the T1 method to both scaffolding proteins as it was used in the previous study of the AKAP79 (21, 46) (Fig. S13, S14). We found that DDC-corrected images still showed significant deviations from simulated random distributions, indicating the presence of clustering. However, the degree of clustering was significantly reduced when compared to the uncorrected and T1-corrected images for both proteins (Fig. 3A). We further confirmed these results at the ensemble level by computationally varying the labeling density of these two proteins using a previously published method (Fig. S15, Supporting Material) (38).

**Figure 3:**
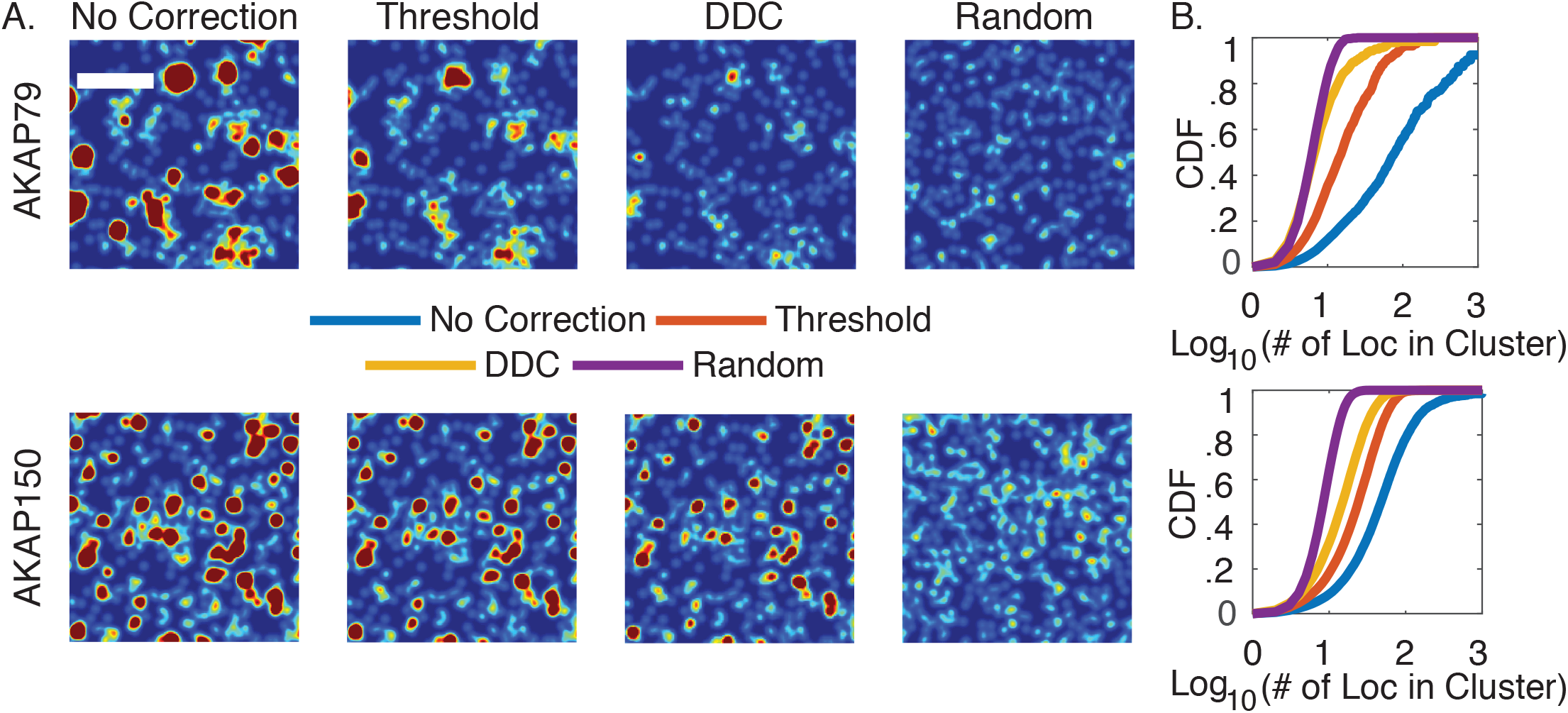
Application of DDC to experimentally measured spatial distributions of AKAP79 and AKAP150. A. SMLM images of the two scaffold proteins without correction, corrected using the thresholding method T1 and DDC, and that of a simulated random distribution using the same number of localizations of DDC-corrected images. B. Cumulative distributions for the number of localizations within each cluster for each protein. (Scale bar, 1*μm*)

To quantitatively compare these images, we used a tree-clustering algorithm (Supporting Material) to group localizations in individual clusters and plotted the corresponding cumulative distributions in Fig. 3B. The cumulative distributions show that interestingly, AKAP150 has a higher degree of clustering when compared to AKAP79, with more than 50% of the localizations within clusters containing greater than 15 localizations, twice that of AKAP79. These results suggest that the clustering of the AKAP scaffolds are differentially regulated and the context dependence is likely important in considering the microdomain-specific signaling functions of the clusters. These accurate, quantitative comparisons of cluster properties would be difficult to achieve by other threshold-based methods.

### DDC identifies both subcellular locations and oligomeric states of dynein

The single molecule nature of SMLM allows one to identify both the subcellular location and copy number of individual molecular components in complexes. However, errors due to repeat localizations lead to misassignment of individual complexes of differential assembly states to incorrect subcellular locations, confounding possible biological interpretations. Previously, using a well-defined DNA origami structure as a calibration standard, SMLM studies showed that dynein, a cytoskeletal motor protein responsible for retrograde transport on microtubules, can exist in monomeric, dimeric, and multimeric states (39). Monomeric dynein was found randomly in the cytoplasm, most likely corresponding to subunits not incorporated into fully assembled motors, which are dimers. Multimeric dynein motors containing two or more dimers were found to arrange into nanoclusters mostly along microtubules, likely involved in coordinated and fine-tuned transport of organelles in the crowded cytoplasm (39). Understanding how dynein motors are arranged inside cells with their respective assembly states provide insight into the function and regulation of dynein in organelle transport. This system also provides a previously quantified experimental system to investigate how blinking-caused artifacts can influence the assignment of individual molecular assemblies.

We performed SMLM imaging on anti-GFP antibody-labeled HeLa IC74 cells that stably express GFP-fused dynein intermediate chain (Fig. 4A, Supporting Material) (39). We then applied the Thresholding method (T1) and DDC to the resulting raw images, with zoomed-in sections shown (white box i) in Fig. 4B. We observed that both the Threshold method and DDC had a lower amount of signal when compared to that from Raw localizations (Fig. 4B, white box i), demonstrating that a significant number of Raw localizations were repeat localizations. Importantly, we also observed that the difference between the threshold- and DDC-corrected images was not constant throughout the images (Fig. 4B, last row), suggesting different assignments of multimeric state for individual dynein assemblies between different methodologies.

**Figure 4:**
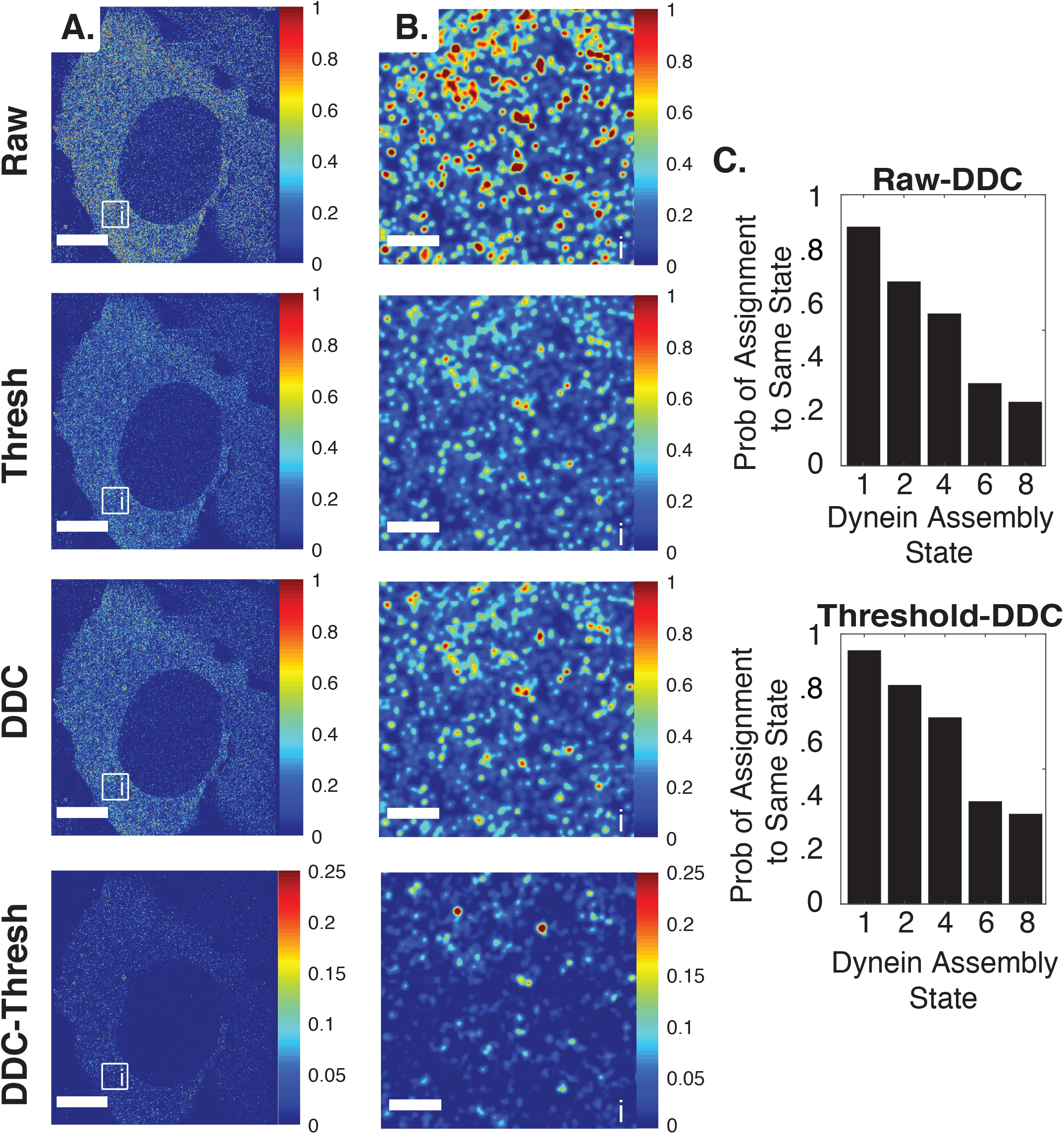
Application of DDC to experimentally measured spatial distributions of dynein. A. SMLM images of dynein for a whole cell with all three method and the difference between the DDC and threshold images (10*μm* scale bar). B. Zoomed in images showing the Raw, Threshold (T1) DDC corrected images and DDC minus Threshold images (1*μm* scale bar). C. The probability of an individual assembly being assigned the same oligomerization state as assigned with DDC for the Raw (top) and Threshold (T1, bottom) methodology (Supporting Material). Note: because a functional dynein motor is homodimeric we only included even number complexes and the monomeric state as previously done in Zanacchi *et al*. (39).

To investigate this difference further, we assigned oligomeric states to individual assemblies from each methodology so that the fractions of each oligomeric state matched what was calibrated in the work of Zanacchi *et al*. (39) (Supporting Material). We then compared the assignment of individual assemblies between the different methodologies by calculating the probability of assigning the same oligomeric state to the same individual complex using two different methods. In Fig. 4C we show that for single dynein monomers both the Raw and Threshold methodologies are in relative agreement with the assignment of DDC (probability > 90%). However, we observed that the higher oligomeric states assigned by both the Raw and Threshold methods had considerable deviations from that of DDC, resulting in different spatial distribution of oligomeric dynein motors in cells. These results demonstrate the importance of using the correct method to obtain both subcellular locations and the quantitative properties of molecular assemblies.

### DDC minimizes measurement noise in labeled symmetric sister chromatin fibers

In addition to quantifying the number of molecules in molecular assemblies and the corresponding sub-cellular locations, DDC can also be applied to minimize noise in the measurement of cellular structural features such as shape and symmetry. To demonstrate such an application, we examined the symmetric structure of sister chromatin fibers. Previous studies have shown that during stem cell differentiation, *Drosophila melanogaster* male germline stem cells undergo asymmetric division to produce a self-renewing stem cell and a differentiating daughter cell (49). The asymmetric division is likely directed by unidirectional replication fork movement and biased histone incorporation between two sister chromatids (40, 50).

To provide a quantitative comparison standard for analyzing DNA and protein contents in sister chromatids, we performed SMLM imaging on YOYO-1 stained chromatin fibers isolated from *Drosophila melanogaster embryos* (Supporting Material, Fig. 5A). For this system the chromatin fibers isolated from embryonic, non-stem cells should exhibit homogenous and symmetric labeling on both sisters. We then applied the threshold (T1) and DDC methods to the raw SMLM images (Fig. 5A). In many fibers we could resolve two parallel sister chromatins; the apparent width of each sister was ≈140 nm (Full width at half maximum, FWHM) and the separation between sisters was ≈200 nm. These characteristics were measured from the projected localizations along the length of fibers. Additionally, DDC-corrected images had a more homogenously distributed signal along the length of chromatin fibers compared to the raw or threshold-corrected images. This scenario is similar to the intersecting filamentous structures with high labeling density presented in Fig. 2.

**Figure 5:**
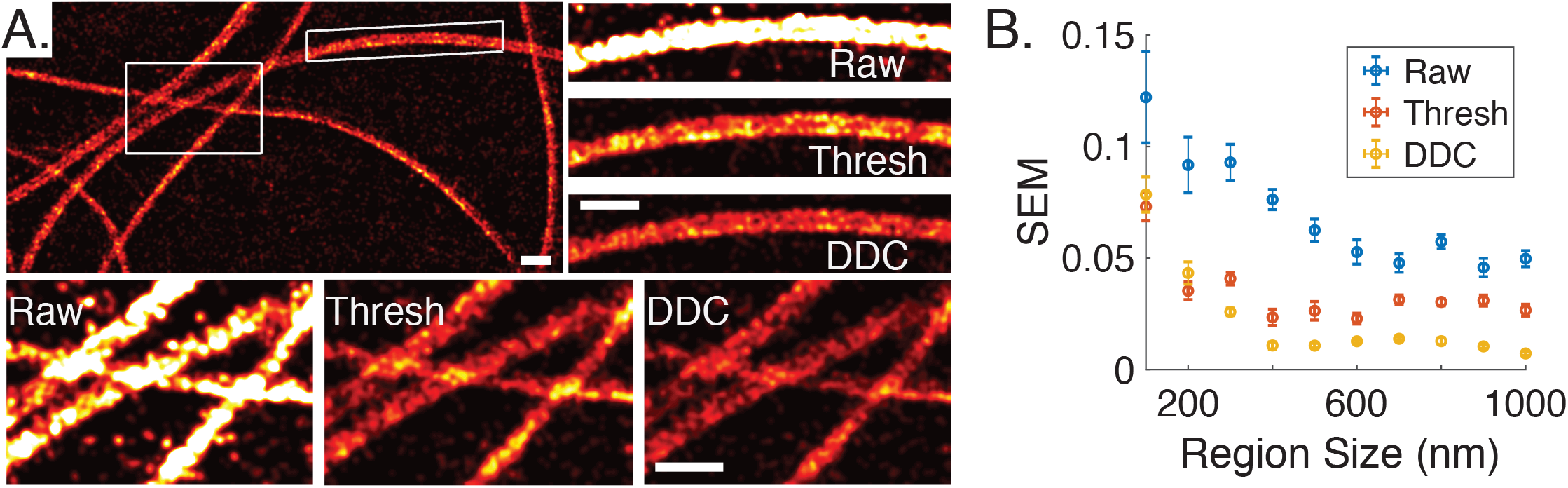
A. Sister chromatids analyzed with DDC and zoom in images showing the resulting images for each of the methodologies (scale bar 1*μm*). B. The standard errors of mean vs. region size for the different methodologies (error bars SEM determined from bootstrapping).

Next, to determine whether the two sister chromatin fibers have a similar amount of DNA, we quantified the ratio of YOYO-1 signal (number of localizations) between the two using segments of different lengths (≈1*μm* was used in the original work of Wooten *et al*. (40) (Supporting Material). Two sisters having identical replicated DNA content would have a ratio of 1 irrespective of the average length of segment used. As shown in Fig. S16, while the ratios of signal between the two sisters for all three methodologies (Raw, Threshold (T1), and DDC) are approximately centered around 1.0, the degree of the ratios’ spread vary considerably, suggesting that while repeat localizations may not affect the accuracy of these measurements, they may instead affect the precision.

To investigate this variation further, we calculated the standard error of the mean (SEM) for the different segment lengths (Fig. 5B). We observed that the SEMs from raw images were the greatest across different segment lengths, and that obtained from DDC were consistently the lowest for segments greater than 300 nm. When the segment lengths became too short, the level of variations became indistinguishable between DDC and the thresholding method due to the intrinsic stochastic labeling density in this experiment. Nevertheless, the apparent SEMs in raw and threshold-corrected images at length scales of chromatin fibers (300 nm to 1 *μ*m) could mask asymmetries in labeled sister chromatin fibers isolated from germ line stem cells (previously quantified with this technique (40)), making it difficult to identify corresponding molecular mechanisms contributing to asymmetry. In summary, this example illustrates how the mishandling of repeat localizations lowers precision and demonstrates the need of DDC when measuring cellular structural features with SMLM.

### Considerations in the application of DDC

As with any method, successful application of DDC to SMLM images requires an understanding of critical factors that could influence the performance of DDC. In this section, we evaluate the impact of localization density and activation rate on the performance of DDC using simulations. We also demonstrate that the commonly used practice of ramping the UV activation power in SMLM imaging should be avoided when applying DDC.

To quantify the influence of localization density on the performance of DDC, we simulated random distributions of fluorophores with different densities ranging from 1000 raw localizations to 15000 localizations per 1*μm*^2^. Note that a density greater than 5000 localizations/*μm*^2^ corresponds to a Nyquist resolution of 30 nm or better. As shown in Fig. 6A, the Image Error increases as the localization density increases and reaches a plateau at ~ .35. We found that the increase in Image Error at high localization densities was mostly due to the decreased raw Image Error of the uncorrected images at high localization densities (Fig. S17A). The decreasing improvement of DDC at increasing sampling rate suggests that a high sampling rate of the underlying structure reduces the image distortion caused by repeats, although very high labeling densities (> 10,000 localizations/*μm*^2^) is usually difficult to achieve for protein assemblies.

**Figure 6:**
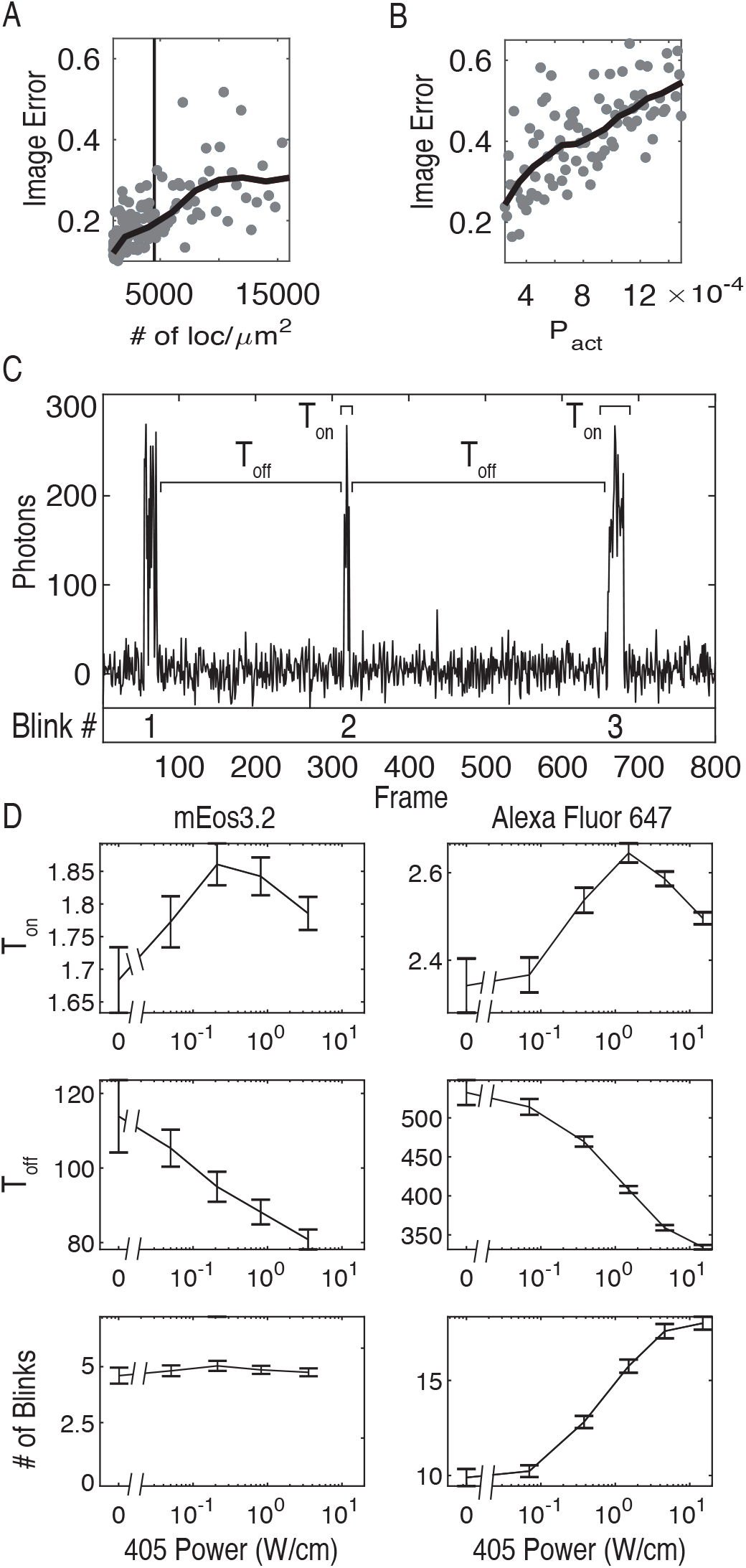
Image Error at different densities of localizations (A) and activation probability per frame (B). The raw data points are shown as gray points and the moving average is shown in black (Supporting Material). C. An intensity trajectory of a single mEos3.2 molecule with labels showing the definitions of *T_on_* and *T_off_*. D. The average *T_on_*, *T_off_*, and number of blinks for Alexa647 and mEos3.2 at different UV activation intensities (405 Power, error bars are standard deviation of mean using two repeats).

Next, to quantify the influence of the activation rate, we varied the activation probability of each simulated fluorophore from .025 to .15 per frame, with 1000 fluorophores randomly distributed throughout a 1*μm*^2^ area. Fig. 6B shows that the Image Error of DDC steadily increases with the activation rate. This increase was because at high activation rates, the temporal overlaps of individual fluorophores that were spatially close to each other increased, which made it difficult to distinguish the repeat localizations from different fluorophores. This trend holds true for all other blinking-artifact correction methodologies. Therefore, as with others, DDC obtains the best images when the activation rate is slow.

Finally, we illustrate one critical requirement for the successful application of DDC, that is, the photokinetics (blinking behavior) of the fluorophore, must be kept constant throughout the acquisition of the SMLM imaging stream (Supporting Material). Note that this requirement is also needed for all other blinking-artifact correction methods (21, 23, 25). One common practice in SMLM imaging is to ramp the activation power gradually throughout the SMLM imaging sequence in order to speed up the acquisition at later times when the number of fluorophores in the view field gradually deplete. The assumption is that activation power only changes the activation rate of a fluorophore (i.e. the probability of a fluorophore being activated per frame), but not the photokinetics of its blinking behavior (i.e. number of blinks, dark time and fluorescence-on time). Such a scenario indeed was shown for the photoactivatable fluorescent protein Dendra (28), but there are also reports showing that the photokinetics of mEos2 and PAmCherry are sensitive to the activation intensity (27, 28).

To illustrate the activation power-dependence of the blinking behaviors of commonly used fluorophores in SMLM, we investigated the photoactivatable fluorescent protein mEos3.2 and the organic fluorophore Alexa647 with different activation (405nm) intensities. We quantified three parameters, number of blinks, off-times (*T_off_*) and on-times (*T_on_*), and reported the mean value for each parameter as a function of activation intensity (Fig. 6C). We define one blink event as one continuous emission event that could span multiple fluorescence on-frames, the number of blinks as the number of repeated emissions separated by dark frames from the same fluorophore, *T_off_* as the time between each blink and *T_on_* as the time that the fluorophore remained fluorescent at each blink-on event (Fig. 6C). We observed that both fluorophores had a similar dependence of *T_on_* with UV intensity, where *T_on_* initially increased and then decreased at higher UV intensities (Fig. 6D, top), suggesting that UV also participates in the fluorescence emission cycle of the fluorophores. Next, we found that *T_off_* decreased non-linearly as the UV intensity increased for both fluorophores (Fig. 6D, middle). Finally, we observed that the average number of blinks for Alexa647 increased dramatically with UV intensity while that of mEos3.2 remained largely constant (Fig. 6D, bottom), suggesting a differential influence of UV in changing the photokinetics of different fluorophores. Thus, varying the activation intensity during the acquisition of a SMLM image can indeed change the blinking characteristics of the fluorophores, which would affect the performance of DDC. These results suggest that changing the activation intensity should only be done when a quantitative approach is not needed, or the proper controls have been performed to show that the fluorophore is insensitive to variations in the activation intensity.

## Discussion

In this work we provided a blinking-artifact correction methodology, DDC, that does not depend upon exact thresholds, additional experiments, or a specific photo-kinetic model of the fluorophore to obtain an accurate reconstruction and quantification of SMLM superresolution images. DDC works by determining a “ground truth” about the underlying organization of fluorophores, the true pairwise distance distribution. We verified by simulations and experiments that such a true pairwise distance distribution can be obtained by taking the distances between localizations that are separated by a frame difference much longer than the average lifetime of the fluorophore. Using the true pairwise distribution, the likelihood can be calculated, where upon maximization of the likelihood one obtains an accurate representation of the true underlying structure.

We compared the performance of DDC with four different thresholding methods using simulated data with various spatial distributions and on fluorophores with different photokinetic models. DDC outperformed these methods by providing the “best” corrected images as well as excellent estimates of the number of molecules in each image. We then experimentally demonstrated that blinking-caused repeat localizations can lead to artificial clustering of membrane scaffolding proteins, misassignment of oligomeric state of dynein at different subcellular locations, and misidentification of DNA content in symmetric sister chromatin fibers. DDC was able to alleviate these artifacts by providing SMLM images devoid of repeat localizations, allowing accurate, quantitative analyses.

Finally, we demonstrated that the higher the activation rate and the density of fluorophores are, the smaller the relative improvement of DDC will be. Note that this applies to all other methods used to eliminate repeat localizations in SMLM imaging. We also showed that in order to use DDC, the common practice of ramping the UV should be avoided in certain cases (depending upon the particular fluorophore), as we verified that mEos3.2 and Alexa647 exhibited activation power-dependent photokinetics. In essence, DDC is best suited for SMLM imaging when quantitative characterizations of heterogenous cellular structures are required. The complete package of DDC is available for download at https://github.com/XiaoLabJHU/DDC. Because of the simplicity and robustness of DDC, we expect it become a field standard in SMLM imaging for the most accurate reconstruction and quantification of SMLM images to date.

## Acknowledgements

Supported by NIH 5T32GM007231 and F31GM115149-01A1 (M.W.), NIGMS/NIH R01GM112008 and R35GM127075, and the Howard Hughes Medical Institute (X.C.).

