## Supporting Material for "A Pairwise Distance Distribution Correction (DDC) algorithm to eliminate blinking-caused artifacts in super-resolution microscopy"

**5 Algorithms 18**

**10 Supporting Material Figures**
Fig. S1: Toy model and justification of main principle
Fig. S2: Photo-kinetic models of fluorophores used in this work
Fig. S3: The convergence of pairwise distance distributions for all simulation systems
Fig. S4: Scatter plots of PALM images used to justify principle

Fig. S5: Illustration showing the calculation of $\mathbf{M}$
Fig. S6: The result of maximizing the likilihood
Fig. S7: Maximization of likelihood results in correct conformation of localizations
Fig. S8: Examples of $\mathbf{M}$ and $P_{R1}(\Delta r|\Delta n)$
Fig. S9: Example of MCMC phase space search
Fig. S10: Resulting error in using methodology of Annibale et al.
Fig. S11: Determining thresholds using methodology of Coltharp et al.
Fig. S12: Results for the 1 dark state fluorophore
Fig. S13: Scatter plots of AKAP79
Fig. S14: Determining thresholds for AKAP79 and AKAP150
Fig. S15: Methodology of Span et al.
Fig. S16 Ratio between sisters
Fig. S17: Raw image error for uncorrected images with varying density and activation rate

### 1 Mathematical justification for true pairwise distance distribution

Here we provide a mathematical justification supporting the principle that the true pairwise distance distribution is obtained when the pairwise distances are taken between localizations separated by a frame difference much longer than the average lifetime of the fluorophore.

Blinking causes the position of a fluorophore to appear throughout multiple frames, we refer to the localizations from the same fluorophore as a blinking trajectory and we define the first localization in a blinking trajectory as the true localization and all subsequent localizations as repeats. An illustration of two blinking fluorophores for a one dimensional image is shown in Fig.S1 with the true localizations of the fluorophores shown as green dots and repeats in red. For this justification we assume that the blinking behavior of the fluorophores are independent of each other and the photo-kinetics of the fluorophores are constant and uniform throughout the acquisition of the image. Note: this is one of the major assumptions needed to apply DDC.

The number of repeats for an arbitrary fluorophore, $a$, follows an unknown random variable, $num_b(a)$, and the determination of the true position of fluorophore $a$, $x_a$, is dependent upon the localization precision of the microscope, $\delta$. For the toy model in Fig.S1 we have no error in determining the position of the fluorophore for simplicity. The distances contributed by two arbitrary fluorophores within an image can then be split into the three arrays/categories below:

$$\left\{ \begin{array}{c} C1 = \sqrt{((x_a+\delta)-(x_b+\delta))^2} \\ C2(1:\gamma) = \sqrt{((x_a+\delta)-(x_b+\delta))^2} \\ C3(1:\gamma') \approx \sqrt{((\delta)-(\delta))^2} \end{array} \right\},$$

where $\gamma = (num_b(a) + num_b(b) + num_b(a) \times num_b(b))$ and $\gamma' = \sum_{n=0}^{num_b(a)} n + \sum_{n=0}^{num_b(b)} n$, are the number of distances contributed to the pairwise distance distribution for the different categories. Here we should note that the number of distances contributed by the repeats [C2 and C3] can be much higher than the distances contributed by the true localizations, $C1$. The pairs of localizations belonging to the three categories for the two fluorophores are shown in Fig.S1 for reference.

The distances in each of the categories are separated in time by a certain number of frames, $\Delta n$. We define $N$ as the maximum lifetime of a fluorophore. The fact that the fluorescent fluorophores have a limited lifetime creates constraints on the frame differences the distances in each category can posses. The possible frame differences for the distances in categories $C2$ and $C3$ are then the following:

$$\left\{ \begin{array}{c} \Delta n_{C1} - N < \Delta n_{C2} < \Delta n_{C1} + N \\ \Delta n_{C3} < N \end{array} \right\},$$

where $\Delta n_{C1}$ is the frame difference between the true localizations in $C1$.

Notice that if we only use the distances between localizations that are separate in time by $N$, $\Delta n = N$, a pair of arbitrary fluorophores that have at least some localizations in their blinking trajectories with a frame difference of $N$ will contribute a certain number of distances, from $C1$ and $C2$ and all of the distances in $C3$ will be eliminated.

Now, if we use the distances with $\Delta n = N$, the number of distances contributed from $C1$ and $C2$ from any pair of arbitrary fluorophores follows the unknown random variable $\phi$. [The distances contributed by each pair of fluorophores follows the same unknown random variable because the photo-kinetics of each fluorophore is the same.]

Then, to obtain an accurate approximation of the true pairwise distance distribution, $P_T(\Delta r)$, we construct the probability distribution with a bin width $\delta$, assume that the pairs of arbitrary fluorophores ($pairs(i)$) within each distance bin $i$ is large, and use the distances between localizations that are separated in time by $N$. The approximate true probability of observing a distance within bin $i$ is then the following:

$$P_d^i(\Delta r|\Delta n = N) = \frac{\sum_{w=1}^{pairs(i)} \phi}{\sum_{q=1}^{All_{pairs}} \phi} \approx \frac{pairs(i) \times \overline{\phi}}{All_{pairs} \times \overline{\phi}} = \frac{pairs(i)}{All_{pairs}} = P_T^i(\Delta r), \tag{1}$$

where $All_{pairs}$ is the number of pairs of fluorophores, $\overline{\phi}$ is the mean of the random variable and $P_d^i(\Delta r|\Delta n = N)$ is the bin $i$ of the pairwise distance distribution between all localizations separated by the given frame. Equation 1 shows that with the previously mentioned assumptions the probability of finding a distance within each bin will be identical to that of the true pairwise distance distribution, justifying the principle. Note that each frame difference larger than $N$ can be used to approximate the true pairwise distance distribution, therefore creating the pairwise distance distribution using all distances between localizations that are separated by a frame difference larger than $N$ leads to an even better approximation of the true pairwise distance distribution.

### 2 The Inner Workings of DDC

#### 2.1 Defining the Likelihood

Here we define the Likelihood as the following:

$$\mathcal{L}(\{R, T\}|\mathbf{r}, \mathbf{n}) = \prod_{i,j \in \{T\}} P_T(\Delta r_{i,j}) \times \prod_{i \in \{R\}, j \in \{R,T\}} P_{R1}(\Delta r_{i,j}|\Delta n_{i,j}), \tag{2}$$

where $\{R, T\}$ are sets that contain the indices of the localizations that are considered the repeats $\{R\}$ and the true localizations $\{T\}$, where both sets account for every localization. The actual experimental data are stored within the two terms $\mathbf{r}$ & $\mathbf{n}$, with the prior containing the coordinates of every localization and the later containing the frame. The first term on the right determines the probability of observing all of the distances between every pair of true localizations. Here the probability distribution $P_T(\Delta r_{i,j})$ is the true pairwise distance distribution, which gives the probability of observing a distance $\Delta r$ between the two localizations $i$ & $j$ if they are both true localizations. The second term is the probability of observing all of the distances between the pairs of localizations if at least one is considered a repeat. Here, the probability distribution $P_{R1}(\Delta r_{i,j}|\Delta n_{i,j})$ gives the probability of observing the distance between the pair of localizations given the frame difference between them if at least one of the localizations is a repeat. Note that every pair of localizations are within the likelihood calculation no matter which localizations are assigned to the sets $\{R \;\&\; T\}$.

Overall, by maximizing the Likelihood a subset of true localizations is determined, where the pairwise distances between the true localizations are independent of frame, $\Delta n$, and follow $P_T(\Delta r)$. Below we provide all additional information needed to calculate $\mathcal{L}(\{R, T\}|\mathbf{r}, \mathbf{n})$. First we discuss how to determine the second distribution $P_{R1}(\Delta r_{i,j}|\Delta n_{i,j})$ and second the methodology for determining the two sets $\{R \;\&\; T\}$.

##### 2.1.1 Determining $P_{R1}(\Delta r|\Delta n)$

To determine $P_{R1}(\Delta r|\Delta n)$ we utilize the pairwise distance distributions between localizations with a given frame $P_d(\Delta r|\Delta n)$ and the true pairwise distance distribution $P_T(\Delta r)$. Here $P_T(\Delta r)$ is known, determined using the pairwise distances between localizations that are separated by a frame greater than $N$ (See Main Text).

Again, the desired distribution $P_{R1}(\Delta r|\Delta n)$ gives the probability of observing a distance between localizations for a given $\Delta n$ if at least one of them is a repeat. $P_{R1}(\Delta r|\Delta n)$ is therefore made up of the distances between $\{R$ and $T\}$ and $\{R$ and $R\}$, where the curly brackets with the *and* indicate the pairwise distances between the localiztions within the sets. While $P_d(\Delta r|\Delta n)$ is made up of the distances between $\{R$ and $T\}$, $\{R$ and $R\}$, and $\{T$ and $T\}$ for a given $\Delta n$. Therefore, $P_{R1}(\Delta r|\Delta n)$ is equal to $P_d(\Delta r|\Delta n)$ with the contribution from the distances between true localizations removed, $\{T$ and $T\}$.

To properly eliminate the part of the distribution that is due to the distances between the true localizations, we quantify the makeup of $P_d(\Delta r|\Delta n)$ and then proportionally remove $P_T(\Delta r)$ from $P_d(\Delta r|\Delta n)$, resulting in $P_{R1}(\Delta r|\Delta n)$.

$P_d(\Delta r|\Delta n)$ is itself a combination of two distributions $P_T(\Delta r)$ & $P_{blink}(\Delta r)$, where the distances between different fluorophores follow $P_T(\Delta r)$ [Categories C1 and C2] and the distances between localizations from the same fluorophore follow $P_{blink}(\Delta r)$ [Category C3]. Here the probability distribution $P_{blink}(\Delta r)$ is the probability of observing a distance between a pair of localizations that are from the same fluorophore [Category C3] and is determined by the resolution of the SMLM experiment.

We can determine $P_{blink}(\Delta r)$ by comparing $P_T(\Delta r)$ to $P_d(\Delta r|\Delta n < N)$. The distribution $P_T(\Delta r)$ by definition lacks all distances between pairs of localizations that are from the same fluorophore and only contains distances between localizations from different fluorophores [Categories C1 and C2]. While $P_d(\Delta r|\Delta n < N)$ not only contains the distances between pairs of localizations from the same fluorophore [Category C3], but the distances between different fluorophores [Categories C1 and C2]. Note that within a SMLM experiment the resolution is very high, and therefore the distances between the localizations from the same fluorophore are very small, much less than 1000 nm. Therefore, the "shape" of the tails of the two distributions $P_T(\Delta r)$ and $P_d(\Delta r|\Delta n < N)$ match each other, as they both only contain the distances between different fluorophores (Data not shown). With this understanding in mind, the distribution $P_{blink}(\Delta r)$ can be obtained by subtracting $P_T(\Delta r)$ from $P_d(\Delta r|\Delta n < N)$ so that the probability of observing a distance greater than 1000 nm is approximately zero, and then normalizing so that the distribution sums to one.

To determine the proportion of each distribution making up $P_d(\Delta r|\Delta n)$, $P_d(\Delta r|\Delta n)$ can be fit to the following equation:

$$X(\Delta n) = Fit[(1-X) \times P_{blink}(\Delta r) + X \times P_T(\Delta r)], \quad (3)$$

where X is between 0 and 1.

The proportion of the distances that follow $P_T(\Delta r)$ come from the distances between $\{T$ and $T\}$ and $\{R$ and $T\}$. We must therefore take this into consideration when determining the proportion of $P_{R1}(\Delta r|\Delta n)$ that follows $P_T(\Delta r)$. To adjust the proportion of the distribution that follows $P_T(\Delta r)$ we calculate the ratio of the number of distances from $\{R$ and $T\}$ relative to the number of distances from $\{T$ and $T\}$ and $\{R$ and $T\}$.

This ratio can be determined by calculating the average number of repeats per fluorophore, $num_b$. $num_b$ can be obtained without having to perform any additional experiments, using the approximate probability that a localization is a repeat (See Approximating the Probability a Localization is a repeat Section of this Supporting Material) and Alg. 1 to obtain a relatively accurate estimation as to the number of blinks per fluorophore. (Note: for this calculation $\kappa(density) = 0$ and $\kappa_2(frame) = 0$, discussed later.) Here we should note that $num_b$ could also be determined by experiment, though these experiments can be difficult and are very sensitive to model specific errors.

The ratio of the number of distances from $\{R$ and $T\}$ relative to the number of distances from $\{R$ and $T\}$ and $\{T$ and $T\}$ is then the following (See Mathematical Justification Section of this Supporting Material):

$$\alpha = \frac{num_b + num_b + num_b * num_b}{1 + num_b + num_b + num_b * num_b} = \frac{\#\{R \text{ and } T\}}{\#\{R \text{ and } T\} + \#\{T \text{ and } T\}}. \quad (4)$$

where $\#\{R$ and $T\}$ indicates the number of distances between the localizations within the two sets. The distribution $P_{R1}(\Delta r|\Delta n)$ is then equal to the following equation:

$$P_{R1}(\Delta r|\Delta n) = Norm[P_T(\Delta r) \times X(\Delta n) \times \alpha + P_{blink}(\Delta r) \times [1 - X(\Delta n)]]. \quad (5)$$

Here $Norm$ indicates that the distribution within the brackets is normalized so that it sums to one. The distribution $P_{R1}(\Delta r|\Delta n)$ is a combination of the two distributions that are from the distances between localizations from different fluorophores ($P_T(\Delta r)$)) and the distances between the localizations from the same fluorophore ($P_{blink}(\Delta r)$). The first term ($P_T(\Delta r) \times X(\Delta n) \times \alpha$), first accounts for the proportion of the distribution $P_d(\Delta r|\Delta n)$ that results from the distances between localizations from different fluorophores and then scales this proportion further with $\alpha$, so that the contribution from the distances between the pairs of true localizations are removed. $P_{R1}(\Delta r|\Delta n)$, for the 1 dark state no clustered simulation is shown in Fig. S8A. As expected, there is a large probability for small distances and small frame differences due to the proportion of distances between the blinks of the same fluorophores being large. Then as the frame difference increases, the proportion of distances between the blinks of the same fluorophores decreases and the distribution converges upon the true pairwise distance distribution, Fig. S8A.

##### 2.1.2 Determining the sets {R} and {T}

To assign a localization to either the {R} set (repeat) or the {T} set (True Localization) DDC uses the following:

$$\{R, T\} = Alg_1[\mathbf{r}, \mathbf{n}, \mathbf{M}, \kappa(density), \kappa_2(frame)]. \quad (6)$$

The sets {R} and {T} are determined within $Algorithm$ 1, which uses the parameters and data within the brackets to assign each localization to one of the two sets. The actual experimental data are stored within the two terms $\mathbf{r}$ & $\mathbf{n}$, where $\mathbf{r}$ contains the coordinates of each localization and $\mathbf{n}$ contains the frame. Here, $\mathbf{M}$ is a matrix that contains the information that is used to determine the probability that a localization is a repeat (See Approximating the Probability a Localization is a repeat Section) and $\kappa(density)$ & $\kappa_2(frame)$ are monotonic functions that are determined within the MCMC. The two functions $\kappa(density)$ & $\kappa_2(frame)$ allow DDC to adjust the probability calculation by taking into consideration the local density of the image and the frame of each localization. These are the two functions that vary during the MCMC to maximize the likelihood, defining the two sets. We discuss the specifics of $\kappa(density)$ & $\kappa_2(frame)$ within the section Alg. 1, Linking Localizations into Trajectories.

#### 2.2 Approximating the probability that a localization is a repeat

Depending upon the number of localizations within a SMLM image, the number of subsets of localizations can be extremely large. To speed up the phase space search and to minimize the likelihood of overfitting DDC calculates the approximate probability that each localization is a repeat (within the blinking trajectory) of a prior localization and only investigates the more likely subsets of localizations using the MCMC approach (Alg. 1, Linking Localizations into Trajectories). Below we discuss how the approximate probability that each localization is a repeat can be determined and then describe Algorithm 1, which defines which localizations are true localizations and which are repeats within DDC.

Here we define the matrix $\mathbf{M}$, which gives the probability that a localization is a repeat of a prior localization given a distance, $\Delta r$, and $\Delta n$ between the localizations.

$$\mathbf{M}(\Delta r \in i, \Delta n) = \frac{P_d^i(\Delta r|\Delta n) - \omega \times P_T^i(\Delta r)}{P_d^i(\Delta r|\Delta n)}, \tag{7}$$

where $P_d^i(\Delta r|\Delta n)$ is the raw probability for the distance between two localizations to be in bin $i$, given that they are separated by $\Delta n$, $P_T^i(\Delta r)$ is the true pairwise distance distribution and $\omega = \frac{\sum_{i>>\sigma} P_d^i(\Delta r|\Delta n)}{\sum_{i>>\sigma} P_T^i(\Delta r)}$, where $\sigma$ is the localization precision of the microscope. Here $\omega$ is a scaling factor used to match the tails of the two distributions, as the distance distributions have a similar shape for $\Delta r >> \sigma$. Fig. S5 illustrates this calculation and the assumption about the tails of the distribution. $\mathbf{M}$, for the 1 dark state no clustered simulation is shown in Fig. S8B, as expected, there are high probabilities with small $\Delta r$ and small $\Delta n$, which get lower as $\Delta r$ and $\Delta n$ increase. $\mathbf{M}$ is the matrix that Alg. 1 uses to link localizations into trajectories.

#### 2.3 Alg. 1, linking localizations into trajectories

Here we describe Alg. 1, which DDC uses to determine which localizations are linked into trajectories using the previously defined $\mathbf{M}$ and $\kappa(density)$ & $\kappa_2(frame)$. (See Approximating the Probability a Localization is a repeat) Note: one could easily modify the algorithm and have it take into consideration more information to determine which localizations belong to each set, but at a computational cost and risk of overfitting.

We wanted our methodology to be able to account for heterogeneous distributions of fluorophores within the same image and to incorporate the "time" dependence for the appearance of localizations. Therefore, one single cutoff probability or threshold was avoided. Instead we made the probability at which localizations are linked together into the same blinking trajectory related to the local density of the image before blinking correction and related to the frame of the localization.

Note: during the maximization of the likelihood for all of the systems within this work we could not simply eliminate localizations without taking into consideration the "probability of repeat", as this led to an extremely large phase space and did not converge within a reasonable amount of time.

Here the reasoning for incorporating the density is this: the more dense a region of an image is the more likely that a true localization could be considered a repeat by chance (based off of the probability calculation, see Alg. 1) and therefore the density of the image needs be taken into consideration. To incorporate the heterogeneity of the image DDC determines the local density of each localization before the blinking correction. To do this DDC calculates the number of raw localizations that are within $2\sigma$ (SMLM resolution) and have a frame difference greater than $N$, for each localization. DDC then defines a monotonically increasing function that is a function of the density, $\kappa(density)$ [Initially $\kappa(density) = 0$]. The flexibility of this function allows DDC to handle heterogeneous distributions of fluorophores by taking into consideration the local density of the image for the probability calculation (See Alg. 1).

Note: the shape of this function is determined during the MCMC approach and is discussed within Alg. 2.

The reasoning to include the frame information within the probability calculation is: because more localizations appear at the beginning of the acquisition of an image when compared to the end of the acquisition, localizations would be more likely to be considered repeats at the beginning of the acquisition than at the end by random chance. (Because fluorophores photo-bleach during the acquisition of a SMLM image.) The time dependence is utilized in a similar manner as the density, where a monotonically decreasing function of the frame of each localization is incorporated into the probability calculation, $\kappa_2(frame)$, see Alg 1.

Note: the shape of this function is also determined during the MCMC approach and is discussed within Alg. 2.

To link localizations into trajectories DDC utilizes Alg. 1. This simple algorithm goes through all localizations and links them into trajectories, starting with the localizations that are most similar in frame. To decide whether or not to link two localizations into the same trajectory [or two trajectories into one] the algorithm used the mean of the "probabilities of blink" of the localizations being considered. DDC calculates the probability of being a blink with the matrix $\mathbf{M}$, and then divides the mean probability by $1 + \kappa(density(ii)) + \kappa_2(frame(ii))$. This takes into account the local density and frame of the localization $ii$. If the probability of the localization [or localizations] is larger than .5, then the localizations are combined into the same trajectory. For each trajectory all localizations but the localization with the smallest frame in each trajectory are then considered blinks.

Note: we should mention that the order in which the localizations in Alg. 1 are arranged does have a small influence on the trajectories that are formed, especially if the activation rate is high. Therefore, DDC also varies the order of the localizations during the MCMC approach to obtain different subsets of true localizations (See Alg. 2 of this Supporting Material for further details.)

Note: we found that not including an algorithm of similar structure to Alg. 1 (takes into account the physical process of fluorophore blinking) either resulted in an extremely slow convergence or got stuck in minimums that deviated from the true image. Therefore, including the information within $\mathbf{M}$ is critical for DDC to converge upon the true image. We should also state that we did not perform an extensive search for alternatives and we do realize that improvements to Alg. 1 could be an area of improvement for DDC in future research.

#### 2.4 Alg. 2, MCMC approach to maximize the likelihood

Here we describe Alg. 2, which DDC uses to maximize the Likelihood and obtain the "correct" subset of true localizations.

Algorithm 2 is a simple Markov Chain Monte Carlo (MCMC) approach that utilizes Alg. 1 in the process. The MCMC approach perturbs three parameters, $\kappa(density)$, $\kappa_2(frame)$ and the order of the localizations to determine the "correct" subset of true localizations. For each step, one of the three previous parameters are modified by a small amount and the likelihood is calculated for the particular subset of true localizations determined by Alg. 1, given those parameters. Alg. 2 then keeps the new parameter and resets the best likelihood if the likelihood is greater than the previous best likelihood or accepts the new parameters if the difference of the likelihood with the old likelihood is greater than a uniform random number. An example of a phase space search is shown in Fig. S9, where the maximization of the likelihood results in the results shown in red.

We found that including the MCMC approach to maximize the log of the likelihood led to significant improvements in the correct number of fluorophores calculated for all systems. Furthermore, for the more heterogeneous distributions of localizations, the Small clusters simulation systems, the MCMC approach led to dramatic improvements in the image error, data not shown. Therefore, the MCMC approach is vital to the successful supplication of DDC even though it is the most computationally expensive step of the methodology.

### 3 Evaluating the three most common threshold methodologies and the absolute best image error from thresholding

Here we investigate the three most common threshold methodologies and compare their results with DDC. We also compare DDC to the absolute best Image Error thresholding can produce. We discuss the results from each of the comparisons here and whenever we reference the 2 dark state systems we are referring to Fig. 2 in the main text and whenever we mention the 1 dark state system we are referencing the results shown in Fig. S12.

#### 3.1 Equations for evaluating the different methods

The image error of each methodology was calculated with the following equation:

$$ImageError = \frac{\sum_{i,j}[Norm(CorrectedImage(i,j)) - Norm(RealImage(i,j))]^2}{\sum_{i,j}[Norm(UncorrectedImage(i,j)) - Norm(RealImage(i,j))]^2}, \quad (8)$$

where $i\&j$ go over all pixels within the images, Norm() indicates that the image is normalized so that the maximum intensity is 1 and the lowest intensity is 0, CorrectedImage is the image that results from a blinking corrected methodology, RealImage is the image that results if an image is generated only using the true localizations and UncorrectedImage is the image with no blinking corrected methodology.
The counting error of each methodology was calculated with the following equation:

$$CountingError = \frac{|Methods\#ofloc - Real\#ofloc|}{Real\#ofloc} \times 100, \quad (9)$$

where $Methods\#ofloc$ is the number of true localizations determined by the methodology and $Real\#ofloc$ is the actual number of true localizations.

#### 3.2 2011, Semi-empirical equation to obtain photo-kinetics (T1)

Perhaps the most famous and most widely used methodology to extract the photo-kinetics and correct for blinking is by utilizing a semi-empirical formula developed in 2011 (1). The parameters from the fit to the semi-empirical formula are also often used with the suggested optimal thresholds from Coltharp et al. (2) with a time threshold equal to 2 times the average dark time of the fluorophore.

For this methodology the distance threshold is often set to 1 pixel (100nm) and the time threshold, $t_d$ (dark time) is varied and the number of localizations at each $t_d$ is quantified. Once the longer $t_d$ is determine the time threshold is often set to approximately 2 times the dark time.

To evaluate the effectiveness of this methodology we applied the methodology to the 1 dark state simulation data for the three different distributions of fluorophores, Fig S10. The semi-empirical formula fit well, but the error in the number of fluorophores and the average dark time was very significant, indicating that the methodology is flawed for systems with more than 1 blink per fluorophore. (The percent error for the extracted parameters is shown in the titles of each subplot.) This previously unknown degree of error is likely due to the small number of simulation systems to which the methodology was applied during the development of the methodology. Though, here we feel that we should state that this previous work was vital for informing the field just how important blinking correction can be.

Considering the large amount of error in the extracted parameters, Fig. S10, we choose to assume that the methodology had perfect knowledge of the characteristic times for the dark states for each simulation system. When comparing the error with the time threshold set to 2 times the known dark time the error in the calculated number of fluorophores improved significantly, Fig. S12. For the two dark state simulation data we set the time threshold equal to 2 times the longer characteristic dark time. The results of applying these thresholds are shown in Fig. 2 in the main text.

When compared to DDC across all molecular distributions and fluorophores DDC outperformed this methodology across every metric. Considering this is the only other methodology that does not require additional experiments to quantify the photo-kinetics of the fluorophore, the experiments here suggest that DDC should be utilized in every situation instead of this methodology.

#### 3.3 2013, Stringent thresholds to eliminate possibility of over-counting (T2)

For the thresholding methodology of Puchner et al. (3), they first characterized the photo-kinetics of the fluorophore and then set an extremely stringent time threshold, so that 99% of blinking dark times would be linked together and a distance threshold equal to 4 times the resolution of the experiment. This methodology was mainly developed to eliminate the possibility of blinking leading to the appearance of clusters, but due to the extreme thresholds this method will deplete the intensity of true clusters.

The results of comparing this thresholding methodology to the 1 dark state simulation systems is shown in Fig. S12. For the Image Error in each of the 3 systems DDC was significantly better than this thresholding methodology. The improvement was especially noticeable for the dense 1 dark state system, as the stringent thresholds are expected to be detrimental to dense clusters. Suggesting that DDC is better at obtaining the true underlying distribution of fluorophores.

Interestingly, this methodology performed especially well for the number of fluorophores in the random and Small clusters 1 dark state systems, but failed for the dense system with a percent error around 15%. When compared to DDC for the number of fluorophores, DDC consistently had a percent error less than 5%. Suggesting that DDC is also a more reliable method under this metric for this fluorophore.

The results for comparing this thresholding methodology with DDC for the 2 dark state simulation systems is shown in Fig. 2 in the main text. Across the board DDC was vastly better than this thresholding methodology for both the Image Error and the error in the number of fluorophores. Suggesting that when the photo-kinetics of the fluorophore are more complicated than a simple 1 dark state DDC is especially beneficial when compared to this methodology. Furthermore, this thresholding methodology requires the characterization of the fluorophore, which wastes valuable time and can be experimentally difficult at times.

##### 3.4 2012, Determining thresholds by knowing the number of fluorophores (T3)

In the methodology developed by Coltharp et al. (2) they characterized the fluorophores to determine the number of blinks per fluorophore to determine the time threshold and the distance threshold. To determine the number of blinks per fluorophore Coltharp et al. utilized a low activation (UV) laser and slowly activated the fluorophores so that individual time traces could be easily extracted. In the last section of the results of the main text we show that this methodology is likely flawed and varying activation intensities change the photo-physics of the fluorophores potentially leading to errors in the number of blinks per fluorophore, Fig. 4. Though, further experiments would be needed for that particular fluorophore. Also, even if the time traces are properly extracted from fluorophores with the same photo-physics the fits to the dark time intervals are error prone and model dependent (2).

Assuming perfect knowledge as to the number of blinks per fluorophore for this methodology, we scanned the number of localizations obtained for each time threshold and distance threshold. The ideal thresholds were determined using the thresholds for the minimal error in the number of localizations at the intersection of the time and distance thresholds. Examples of this phase space search for six different systems investigated in this work are shown in the first column of Fig. S11, with the corresponding Image Error for each set of thresholds shown in the second column. (Note: the error is log scale for the first column so one can clearly see why the exact thresholds were chosen.) The thresholds determined by this methodology are shown in the following table:

| System | Time Threshold ($n$) | Distance Threshold (nm) |
|---|---|---|
| Random 1 dark | 25 | 130 |
| Small clusters 1 dark | 20 | 130 |
| Dense 1 dark | 20 | 100 |
| Random 2 dark | 35 | 130 |
| Small clusters 2 dark | 35 | 130 |
| Dense 2 dark | 30 | 100 |
| Filament | 35 | 80 |

Note: Logically, the optimal thresholds for this methodology became less intense the more dense the molecular distributions became.

The results of applying this methodology are shown in Fig. S12 for the 1 dark state systems and Fig. 2 for the 2 dark state systems. Considering with this methodology we assumed perfect knowledge for the number of blinks per fluorophore it was of little surprise that the error in the calculated number of fluorophores was actually lower than DDC for the 1 dark state systems. The error in the number of fluorophores was less than 6% for both methods for all systems for the 1 dark state fluorophore. Even though the error in the number of fluorophores for both methodologies was comparable, the DDC Image Error was lower for each 1 dark state system when compared to this thresholding methodology. Suggesting, that DDC captures a more reliable representation of the true localizations, while resulting in a comparable error in the number of fluorophores for the simple 1 dark state fluorophore.

This was also the case for the 2 dark state simulation systems except for the dense distribution system. For the dense distribution system the error in the number of fluorophores was significantly worse for DDC, about 12%, while the thresholding methodology performed well with this metric. (We should note again that this is assuming perfect knowledge as to the number of blinks per fluorophore, so it is expected that the error in the number of fluorophores will always be low with this methodology.) Even though DDC performed worse for the dense 2 dark state system for the number of fluorophores, for the Image Error DDC greatly surpassed this thresholding methodology for all three distributions of fluorophores. The most significant improvement was for the dense system, where this thresholding methodology performed much worse than even an uncorrected SMLM image. Suggesting that DDC is vastly superior than this thresholding methodology for a more complicated 2 dark state fluorophore and great care should be taken when utilizing this methodology when actual clustering exists.

#### 3.5 The absolute best thresholds for the image error (T4)

Considering DDC was able to surpass all of the traditional thresholding methodologies with regards to the Image Error, we wanted to see if any thresholds could surpass DDC. To do this we scanned the time threshold and distance threshold for each system and picked the thresholds that resulted in the mean minimum Image Error for each of the seven systems. The thresholds picked by this methodology are shown in the following table:

| System | Time Threshold ($n$) | Distance Threshold (nm) |
|---|---|---|
| Random 1 dark | 17 | 160 |
| Small clusters 1 dark | 13 | 170 |
| Dense 1 dark | 5 | 190 |
| Random 2 dark | 39 | 140 |
| Small clusters 2 dark | 28 | 150 |
| Dense 2 dark | 3 | 210 |
| Filament | 43 | 80 |
| Continuous Filaments | 10 | 80 |

The results of comparing the absolute best threshold methodology with DDC is shown in Fig. S12 for the 1 dark state system. As expected this thresholding methodology performed best for the metric of Image Error when compared to the other thresholding methodologies. Interestingly, DDC was still able to outperform the thresholding methodology in terms of the Image Error and in terms of the number of fluorophores.

The results of comparing this thresholding methodology with DDC for the 2 dark state system is shown in Fig. 2. Interestingly, for this fluorophore the Image Error and the error in the number of fluorophores for the Random and the Small clusters systems was similar between the two methods. The major difference was for the dense system where the error in the number of fluorophores was around 80% for the thresholding system, while DDC maintained an error of about 12%. Suggesting that the Image Error for the 2 dark state systems was similar between the two methods, but DDC was able to surpass this thresholding methodology in terms of determining the proper number of fluorophores.

These results suggest that even with the absolute best thresholds DDC is still a more reliable approach in regards to the two metrics investigated within this work.

### 4 Methods

#### 4.1 Methodology of Sphan et al.

The implementation of Sphan et al. was done by randomly selecting subsets of localizations (with replacement) and then using the threshold of 2.5 (just as in (4)) as the definition of a cluster — to create the cluster masks. The normalized average density within the clusters ($P/P_o$) vs. the relative area of the image the clusters covered ($\eta$) was plotted for all subsets of localizations to determine if clustering was significant for the system of interest. For this methodology, clustering is deemed significant if $P/P_o$ rises above 1 and stays above 1.

We tested this method on three different simulation systems (Random, Small Clusters, Dense Clusters) with the two-state fluorophore and show these results in Fig. S15A. We observed that the randomly distributed fluorophores maintained a $P/P_o$ equal to 1 while the Dense cluster system rose significantly well above 1, demonstrating that the methodology could adequately recognize that there were clusters in the Dense cluster system and that there were not clusters in the Random system. As expected an intermediate value for the Small cluster system was also observed.

Next, to investigate the clustering of AKAP79/150 with an orthogonal method to DDC, we applied the methodology of Spahn et al. on the superresolution data of each of the two orthologs. The results of this analysis are shown in Fig. S15B, where $P/P_o$ for both rose slightly above a $P/P_o = 1$. These results support the previous findings that the two are significantly clustered, supporting the analysis as quantified by DDC. Though, we should note that $P/P_o$ did not reach high values (like that for the Dense cluster system), suggesting that just as with DDC, the clustering of the two orthologs are not "extreme."

#### 4.2 Specifics for simulations

First, six different sets of data were simulated, 3 different underlying structures and 2 different fluorophores. The two fluorophores followed the two models in Fig.S2. In these simulations the fluorophores only registered as a localization if it was in the active state. For the different simulations the first condition contained no clusters [Random] and all fluorophores were randomly distributed within a 1000nm by 1000nm square and allowed to blink according to the kinetic models in Fig.S2. The second [Small clusters] and third [Dense] conditions had 3 clusters each with 10% of the fluorophores distributed into the clusters for the Small clusters system and 50% for the Dense system. For each of the simulations with clusters each cluster's central location was randomly defined and the localizations within each cluster followed a normal distribution around that center with a $\sigma = 40$. For each of the six systems 24 different images were generated and analyzed for each methodology.

Second, for the simulations involving filaments, we randomly distributed 50% of the true localizations along 5 lines and randomly deposited the rest. We simulated 24 images, with 1000 true localizations each, with approximately 4000 localizations total, following the photo-kinetic model in Fig.S2A. These simulations produced filaments that were clearly visible, but not homogeneous along the filaments.

Third, to produce "intersecting" continuous overlapping filaments we simulated filaments with no varying label density and with a localization error of 20 nm. This was done by placing a fluorophore every 5 nm along a filament. These simulations also followed Fig.S2A and resulted in images like that in Fig. 2 far right.

#### 4.3 Methods for experiments that were used to calculate $Z(\Delta n)$

##### 4.3.1 Strains

The strains with chromosomal fluorescent protein fusion tags were constructed using $\lambda$-RED-mediated homologous recombination (5). Some results used in this paper came from strains that also harbor a single chromosomal DNA site marker (tetO6), the DNA markers are positioned in various positions on the chromosome, and a portion of the results are not relevant and thus not discussed in this publication. The details for the construction of these bacterial strains are described in detail in a previous publication (5).

##### 4.3.2 Cell growth

For live cell imaging, single colonies were picked from LB plates and cultured overnight in EZ Rich Defined Media (EZRDM, Teknova) with 0.4% glucose, at room temperature (RT) with shaking. The next morning, cells were reinoculated into fresh EZRDM with 0.4% glucose and grown at RT until they have reached mid-log phase (O.D.600 0.3-0.4). For simultaneous visualization of DNA site markers (results are not reported here), cells were harvested and resuspended in fresh EZRDM supplemented with 0.3% L-arabinose and 0.4% glycerol and allowed to grow for two additional hours, these cells were harvested via centrifugation and imaged immediately.

For fixed cell experiments, cells were grown accordingly and fixed in 3.7% (v/v) paraformaldehyde (16% Paraformaldehyde, EM Grade, EMS) for 15 min at RT, washed with 1X PBS and imaged immediately.

##### 4.3.3 Nascent rRNA labeling (smFISH)

We performed smFISH using a previously published protocol ((6), (7)). Briefly, cells were grown in EZRDM glucose as previously described; 5 ml of mid-log phase cells were fixed with 3.7% (v/v) paraformaldehyde (16% Paraformaldehyde, EM Grade, EMS), placed for 30 min on ice. Next, cells were harvested via centrifugation, and subsequently washed two times in 1X PBS. Cells were then permeabilized by resuspending in a mixture of 300 l of H2O and 700 $\mu$l of 100% ethanol and incubating with rotation at RT for 30 min. Cells were stored at 4 °C until next day. Wash buffer was freshly prepared with 40% formamide and 2x SSC and put on ice. Cells were spun-down in a bench-top centrifuge at 10000 rpm for 3 min and the cell pellet was resuspended in 1 ml of wash buffer. The sample was placed on a nutator to mix for 5 min at RT. Hybridization solution was prepared with 40% formamide and 2x SSC, subsequently, dye-labeled oligo probes were added to hybridization solution to a final concentration of 1 $\mu$ M. Cell were spun-down again and 50 $\mu$l of hybridization solution with probe was added to the pellet. The hybridization sample was mixed well and placed overnight in a 30 °C incubator. Next day, 10 $\mu$l of hybridization sample was washed with 200 $\mu$l of fresh wash buffer and incubated at 30 °C for 30 min, this was repeated one more time. The washed sample was imaged immediately: without STORM imaging buffer for ensemble fluorescence, with STORM buffer to induce dye blinking for superresolution imaging. glucose oxidase + thiol STORM buffer was used to image samples with only dye labeling (50 mM Tris (pH 8.0), 10 mM NaCl, 0.5 mg ml-1 glucose oxidase (Sigma-Aldrich), 40 g ml-1 catalase (Roche), 10% (w/v) glucose and 10 mM MEA (Fluka))((8)). Thiol only STORM buffer (10 mM MEA, 50 mM Tris (pH 8.0), 10 mM NaCl) was used to image samples with both endogenously expressed fluorescent proteins and dye labeling. This was to preserve the fluorescent signal from fluorescent proteins, since the presence of glucose oxidase in the STORM buffer tended to quench the fluorescent protein signal. Pre-rRNA transcripts were detected with a single probe L1, conjugated at the 5' with either Alexa Fluor 488 (NHS ester) or Alexa Fluor 647 (NHS ester) (IDT) ((9)). Upon receiving the commercial oligos, a working stock (50 M) was made and aliquoted for storage at -20 °C.

###### 4.3.4 Cell imaging and SMLM analysis

A 3% gel pad made with low-melting agarose (SeaPlaque, Lonza) in EZRDM was prepared. Live cells of an optimal imaging density were deposited onto the gel pad and immobilized with a coverslip for imaging as previously described ((6)). An Olympus IX-81 inverted microscope with a 100X oil objective (UPlanApo, N = 1.4x) was used, with 1.6x additional amplification. Images were captured with an Ixon DU-895 (Andor) EM-CCD with a 13 $\mu$m pixel size using MetaMorph (Molecular Devices). Illuminations (405 nm, 488 nm, 561 nm, 647 nm) were provided by solid-state lasers Coherent OBIS-405, Coherent OBIS-488, Coherent Sapphire-561, and Coherent OBIS-647 respectively. Fluorescence was split using a multi dichroic filter (ZT 405/488/561/647rpc, Chroma), and the far-red, red and green channels were further selected using HQ705/55, HQ600/50 and ET525/50 bandpass filters (Chroma). Gold fiducial beads (50 nm, Microspheres-Nanospheres, Mahopac, NY) were used to correct for any sample drift during imaging. All superresolution images were acquired with a 10 ms exposure time with 3000-9000 frames. Activation of fluorescent proteins was done simultaneously to fluorophore excitation, and activation laser (405) was kept at a constant power throughout the imaging session. For two-color imaging, the simultaneous, multi-color acquisition was achieved using Optosplit II or Optosplit III (Cairn Research), colored channels were overlaid using calibration images from TetraSpeck beads (Life Technologies, T-7279), as previously described ((10)). Initial fitting of raw imaging data was performed via thunderSTORM plugin ((11)). Later analysis of localizations with DDC was processed using custom Matlab scripts, which will be made available upon request.

##### 4.4 Methods used for sister chromatid experiments

###### 4.4.1 Chromatin fiber preparation from Drosophila melanogaster embryos with YOYO-1 staining:

Young embryos (<2 hours old, 15-20 embryos per experiment) were collected and washed 3 times in room temperature lysis buffer (100mM NaCl, 25mM Tris-base, 0.2% Joy detergent, pH 10; adapted from McKnight and Miller (12)). Embryos were transferred to the center of a clean glass slide (Fisherbrand Superfrost Plus Microscope Slides) and subsequently drained of residual lysis buffer. Following removal of residual lysis buffer, 20 $\mu$l of fresh lysis buffer was then added to the surface of the glass slide to immerse embryos. Embryos were then manually broken apart with dissecting forceps to release embryonic nuclei from the intact embryo. After breaking open the embryo, the protective outer layers of the embryo (chorion layers, waxy layer and vitelline membrane) were removed, and the nuclei were allowed to sit in lysis buffer until fully lysed ( 2 minutes). 10 $\mu$l of sucrose/formalin solution (1M sucrose; 10% formaldehyde) was then added on top of the lysed nuclei, after which, a large coverslip (22x50mm; Thermo Scientific? Rectangular Cover Slips) was placed on top of the lysed chromatin solution and incubated for two minutes at room temperature. Following incubation, slides containing chromatin fibers derived from lysed embryonic nuclei were transferred to liquid nitrogen and allowed to sit for two minutes. Slides were then removed from liquid nitrogen, after which, the cover slip was removed with a razor blade. Slides were then transferred to cold (-20°C) 95% ethanol and incubated for 10 minutes. Slides were removed from ethanol and placed at a 45 deg angle for 2 minutes (or until almost all ethanol has evaporated from the slide, but it is not completely dry). 500 $\mu$l of fixative solution [0.5% formaldehyde in 1xPBST (1xPBS with 0.1% Triton)] was then slowly added to the surface of the slide, after which, the slide was incubated for 2 minutes. Slides were then drained of fixative solution and transferred to a coplin jar containing 50 ml of 1xPBS. To fully wash chromatin fiber samples, slides were then removed from coplin jar and drained of remaining 1xPBS. Used PBS in the coplin jar was then discarded, and the coplin jar was refilled with 50 ml fresh PBS. Slides were then placed back inside coplin jar and incubated at room temperature for two minutes. Slides were removed from coplin jar and placed in fresh PBS two additional times in order to complete the wash process. Following washing, slides were transferred to a humid, dark place and pre-blocked with 500 $\mu$l of blocking solution (2% BSA, MilliporeSigma Bovine Serum Albumin, in 1xPBS) for 30 minutes.

Blocking solution was then drained and 500 $\mu$l of DNA labeling solution containing 1uM YOYO-1-DNA dye (ThermoFisher Scientific Invitrogen YOYO-1) was then slowly added to the surface of the slide. Slides were then incubated for 120 minutes in a humid, dark place. Following incubation, slides were drained of DNA labeling solution and transferred to a coplin jar containing 50 ml of 1xPBS. Slides were removed from coplin jar and placed in fresh PBS two additional times in order to complete the wash process. Following washing, slides were removed from coplin jar and drained of residual 1xPBS. Slides were then mounted in preparation for STORM imaging.

##### 4.4.2 SMLM Imaging

The single molecule localization microscopy (SMLM) imaging of DNA fibers is based on the DNA intercalating dye method (Flors, 2009, PMID: 19554598). The fibers on cover slides were labeled with 1uM YOYO-1 for 120 min. 8-10 $\mu$L dSTORM buffer (Nahidiazar, 2016, PMC4938622) were added on the top of the fibers and sandwiched with a clean coverglass (#1 Fisher Scientific). The coverglass was then sealed with nail polish. The sample can be imaged within 4-5 hours with reasonable localizations. Image acquisitions were performed on an Olympus IX-71 inverted microscope with a 1.49 NA 100 X TIRF objective, a ZT405/488/561 dichroic mirror (Chroma), an ET525/50 emission filter (Chroma), and an Andor iXon Ultra 897 EmCCD camera. Ten to thirty 3000-frame acquisitions of YOYO-1 signal were then obtained with a 30 frame/second rate at 1 kW/cm2 488nm laser power. During the imaging, the activation 405 nm laser was ramped stepwise (Images were analyzed individually and then recombined) up by 1 W/$cm^2$ per movies (3000 frames) to obtain more localizations. dSTORM data were first localized using 2D gaussian fitting in an ImageJ plug-in, ThunderSTORM. A bandpass filter (70 500nm) for sigma was applied to remove the single pixel noise and out-of-focus molecules. The cross-correlation method in ThunderSTORM was applied to correct the long-time scale drift.

##### 4.4.3 Analysis

To quantify the number of localizations between sister chromatids we first fit a spline function to cropped out regions that showed single filaments. We then projected the localizations along this new axis — so that there was no curvature within the filaments and they were centered. We then split the filament into as many specifically sized segments as possible (as varied within the corresponding figures) and quantified the number of localizations in the upper sister relative to the lower sister for the different blinking-artifact methods.

#### 4.5 Methods used for dynein experiments

##### 4.5.1 Cell line

Stably transfected HeLa IC74-mfGFP cells (The dynein intermediate chain is GFP labeled, from Takashi Murayama lab, Juntendo University School of Medicine, Tokyo, Japan) were plated on a 8-well Labtek 1 coverglass chamber (Nunc). Cells were cultured under standard conditions (DMEM, high glucose, pyruvate, 10% FBS and 2 mM glutamine).

##### 4.5.2 Immunostaining

Cells were fixed with PFA (4% in PBS) at RT for 20' and incubated with blocking buffer (3% (wt/vol) BSA (Sigma) in PBS and 0.2% Tryton X-100 (Thermo Fisher Scientific) for 1 hr. Dynein intermediate chain-GFP was immunostained with primary antibody (chicken polyclonal anti GFP, Abcam 13970) diluted 1:500 in blocking buffer for 45 minutes at RT. Cells were rinsed 3 times in blocking buffer and incubated for 45 minutes in secondary antibodies donkey-anti chicken labeled with photoactivatable dye pairs for STORM (Alexa Fluor 405-Alexa Fluor 647).

##### 4.5.3 Imaging

Imaging was done using Nanoimager-S microscope (Oxford Nanoimaging) with the following specifications: 405, 488, 561, and 640 nm lasers, and 665–705 nm band-pass filters, 100× 1.4 NA objective (Olympus), and a Hamamatsu Flash 4 V3 sCMOS camera. Localization microscopy images were acquired with 16-ms exposure and 50,000 frames. 405-nm activation was kept constant and then processed using the NimOS localization software (Oxford Nanoimaging).

##### 4.5.4 Analysis

To quantify each "cluster" as a particular oligomerization state we first quantified the number of localizations within each individual cluster using the hierarchical tree clustering algorithm built within matlab. We then assigned the oligomer state of dynein (for each method) so that the fractions of each state were the same as in (13). We then compared the assigned state for each individual "cluster" as in the main text.

#### 4.6 Methods used for AKAP150

For fixed-cell stochastic optical reconstruction microscopy (STORM) imaging, cells were fixed with 4% paraformaldehyde (PFA) for 20 min and then washed with 100 mM glycine in Hanks balanced salt solution (HBSS) to quench the free PFA. Cells were permeabilized and blocked in a permeabilization solution with 0.1% Triton X-100, 0.2% bovine serum albumin, 5% goat serum, and 0.01% sodium azide in HBSS. The cells were then incubated overnight at 4°C with an anti- AKAP150 (Millipore Sigma 07-210, Cat. # 07-210 EMD Millipore) antibody at a 1:500 dilution, followed by 1 to 2 hours with goat anti-rabbit Alexa 647?conjugated antibodies at 1:1000 dilution. The cells were then post-fixed again in 4% PFA, quenched with 100 mM glycine in HBSS, and washed with HBSS to prepare for imaging. Immediately before imaging, the medium was changed to STORM-compatible buffer [50 mM tris-HCl (pH 8.0), 10 mM NaCl, and 10% glucose) with glucose oxidase (560 mg/ml), catalase (170 mg/ml), and mercapto-ethylamide (7.7 mg/ml). STORM images were obtained using a Nikon Ti total internal reflection fluorescence (TIRF) microscope with N-STORM, an Andor IXON3 Ultra DU897 EMCCD, and a 100x oil immersion TIRF objective. Photoactivation was driven by a Coherent 405-nm laser, while excitation was driven with a Coherent 647-nm laser. Puncta localization was performed using both Nikon Elements analysis software.

#### 4.7 Methods used for characterizing blinking

##### 4.7.1 Sample preparation:

Plac::mEos3.2 plasmid (pXY329) was constructed based on pJL005 (Plac::FtsZwt-mEos3.2) (14) using In-fusion cloning (Takara) to remove the ftsz gene. MG1655 cells were transformed with pXY329 and grow up in M9+ media. The cells are harvested at log-phase and fixed by 3.8% para-formaldehyde in 1X PBS buffer. The fixed cells were washed by 1X PBS for 3 times and saved in 4°C no longer than one week.

Streptavidin conjugated with $AlexaFluor^{TM}$647 (SA-AF647) was purchased from Thermo Fisher Scientific. The SA-AF647 working solution was made freshly every time by diluting original stock ( $36\mu$M) to 10 pM in 1X PBS with 0.5% Tween20.

##### 4.7.2 Imaging

**PALM**: Fixed MG1655-Plac::mEos3.2 cells were sandwiched between a 3% PBS agar-pad and a coverglass as previously described (15). PALM imaging was preformed as previous study (14) on an Olympus IX71 inverted microscope with a 100X, 1.49 NA oil-immersion objective. The 561nm excitation laser power was tuned to 1500 W/cm2 while the 405nm laser power varied from 0 to 3.5 W/cm2. For the 0 W/cm2 condition, a short pulse (1 second) of 3.5 W/cm2 405nm laser was applied to activate some mEos3.2 molecules to red fluorescent state. At each 405-power condition, 6 movies of 3000 frame images with 10ms exposure time were collected continuously. Three repeats of all the 405-conditions were performed to get the average blinking behavior.

**dSTORM**: 10pM SA-AF647 was flown into a preassembled chamber with biotin-PEG coated coverglasses from X for 5min and washed three times with 1X PBS. The STORM buffer was made freshly using the recipe described in (16) and injected to the chamber to replace the PBS buffer before imaging. All STORM images were taken after 60 min since the oxygen level in the buffer was shown to be stable after 1 hour. dSTORM imaging was performed on an Olympus IX81 inverted microscope with a 100X, 1.45 NA oil-immersion objective. The 647nm excitation laser power was tuned to 1800 W/cm2 while the 405nm laser power varied from 0 to 13.9 W/cm2. At each 405-condition, 2-3 5000-frame movies at different regions on the coverglass were taken with a 30ms exposure time. Two repeats of all the 405-conditions were performed.

##### 4.7.3 Data processing

The single fluorophore spots in both PALM and dSTORM movies were localized by an ImageJ (17) plugin ThunderSTORM (18). All the spots with irregular properties (abnormal sigma, too high or low intensity, or multiple spots within 500 nm range) were removed. A customized Matlab code was used to link the same spots within 3-4 folds of localization limitation (100nm) throughout the whole movie using a nearest neighbor algorithm. The continuous frames with localization from the same linked fluorophore were counted as on frames. Other frames before the last on-frame were counted as off frames. Blinking number was calculated as the sum of on frame number.

### 5 Algorithms

**Algorithm 1**

**procedure** DETERMINE WHICH LOCALIZATIONS ARE BLINKS
  $\mathbf{M}(\Delta r, \Delta n) \leftarrow$ Probability that a localization is a repeat of the preceding localization given the Distance and Frame between the preceding localization
  $traj(i) \leftarrow$ is the trajectory that localization $i$ is assigned (before the for loops each localization is assigned it's own personal trajectory)
  $\Delta\mathbf{r}_{traj(i),traj(ii)}$ and $\Delta\mathbf{n}_{traj(i),traj(ii)} \leftarrow$ arrays containing the pairwise distances and frame differences between all localizations in the two trajectories containing localization(i) and localization(ii)
  $\Gamma = length(\Delta\mathbf{r}_{traj(i),traj(ii)})$
  $\kappa(density(i)) \leftarrow$ a monotonically increasing function that is dependent upon the local density of localization(i) without blinking correction (Supporting Material).
  $\kappa_2(frame(i)) \leftarrow$ a monotonically decreasing function that is dependent upon the frame of localization(i) (Supporting Material).
  {T}=1:length(Localizations) $\leftarrow$ the indices that are the true localizations
  {R}=empty array $\leftarrow$ the indices of the localizations that are repeats
  **for** $\Delta$n=1:max(frame) **do**
    **for** i=1:length(Localizations) **do**
      **for** ii={T} **do**
        **if** frame(ii)-frame(i)=$\Delta$n **then**
          **if** $\frac{[\sum_j^{\Gamma} M(\Delta\mathbf{r}_{traj(i),traj(ii)}(j), \Delta\mathbf{n}_{traj(i),traj(ii)}(j))]/\Gamma}{1+\kappa(density(ii)+\kappa_2(frame(ii))} > .5$ **then**
            Combine all the Localizations within the two trajectories into a single trajectory
            Eliminate Localization(ii) from {T} as it is now considered a repeat
            Include Localization(ii) in {R} as it is now considered a repeat

**Algorithm 2**

**procedure** MARKOV CHAIN MONTE CARLO TO MAXIMIZE LIKELIHOOD
  Max Lik=$-\infty$
  Count=1
  Number of Steps=1000
  **while** Count<Number of Steps **do**
    $\kappa(density(:)) = \kappa^{Stored}(density(:))$
    $\kappa_2(frame(:)) = \kappa_2^{Stored}(frame(:))$
    $C = rand \leftarrow$ a random uniform number
    **if** $C < 1/3$ **then**
      Adjust the function $\kappa(density(:))$ by a small amount
      Ensure that $\kappa(density(:))$ is still a monotonically increasing function of density
      Ensure that the mean of $\kappa(density(:))$ over all density values from all localizations equals zero
    **else if** $C < 2/3$ **then**
      Adjust the function $\kappa_2(frame(:))$ by a small amount
      Ensure that $\kappa_2(frame(:))$ is still a monotonically decreasing function of the frame
      Ensure that the mean of $\kappa_2(frame(:))$ over all localizations equals zero
    **else**
      Perturb the order of localizations that have the same frame ▷ This will change which localizations are linked together into the same trajectory
    $\{R, T\} \leftarrow$ Perform Alg. 1 with new $\kappa(density(:))$, new $\kappa_2(frame(:))$, and in new defined order
    Lik $\leftarrow$ Calculate log likelihood with new Corrected Localizations
    **if** Lik>Max Lik or $log(rand) < |MaxLik - Lik|$ **then**
      Store new parameters
      Max Lik=Lik
    **else**
      Go back to old parameters
    Count=Count+1

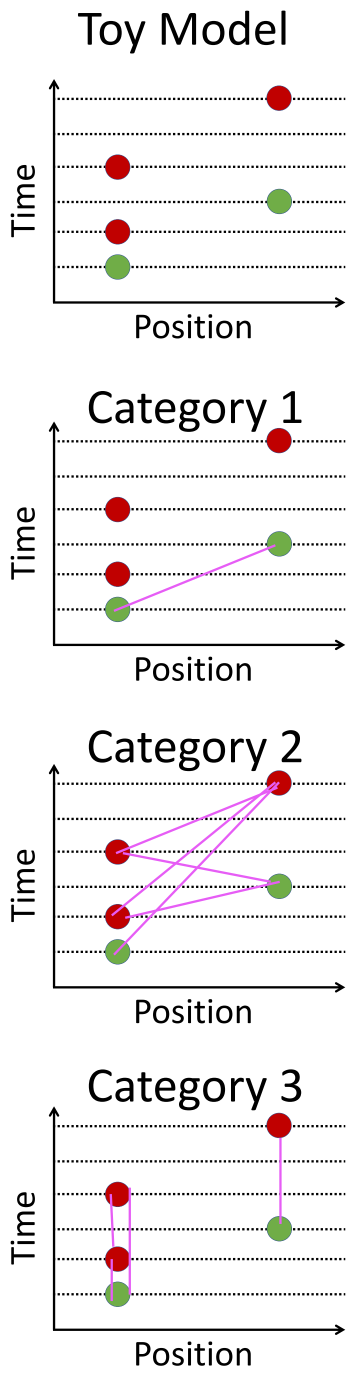

Figure S1: The top row shows a simple one dimensional system illustrating the blinking of two fluorophores, where the green dots are the true localizations and the red dots are repeats. The subsequent rows show the different categories referenced within the Supporting Material, with the pink lines illustrating the pairs of localizations for each category.

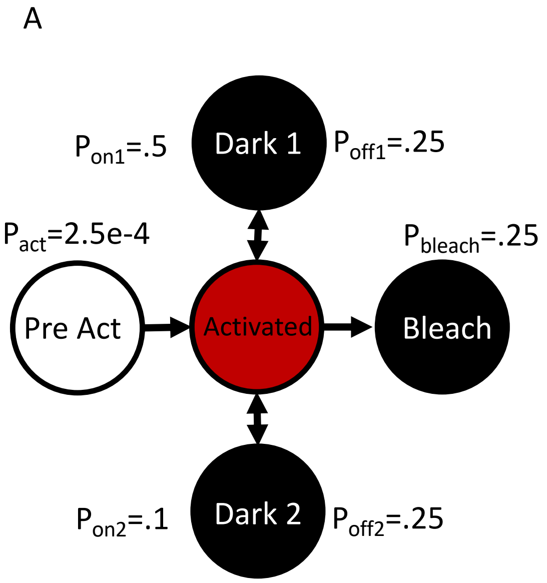

Figure S2: The two kinetic models used to simulate blinking, A.) 2 dark state and B.) 1 dark state. The transition probabilities per frame are shown in the figure.

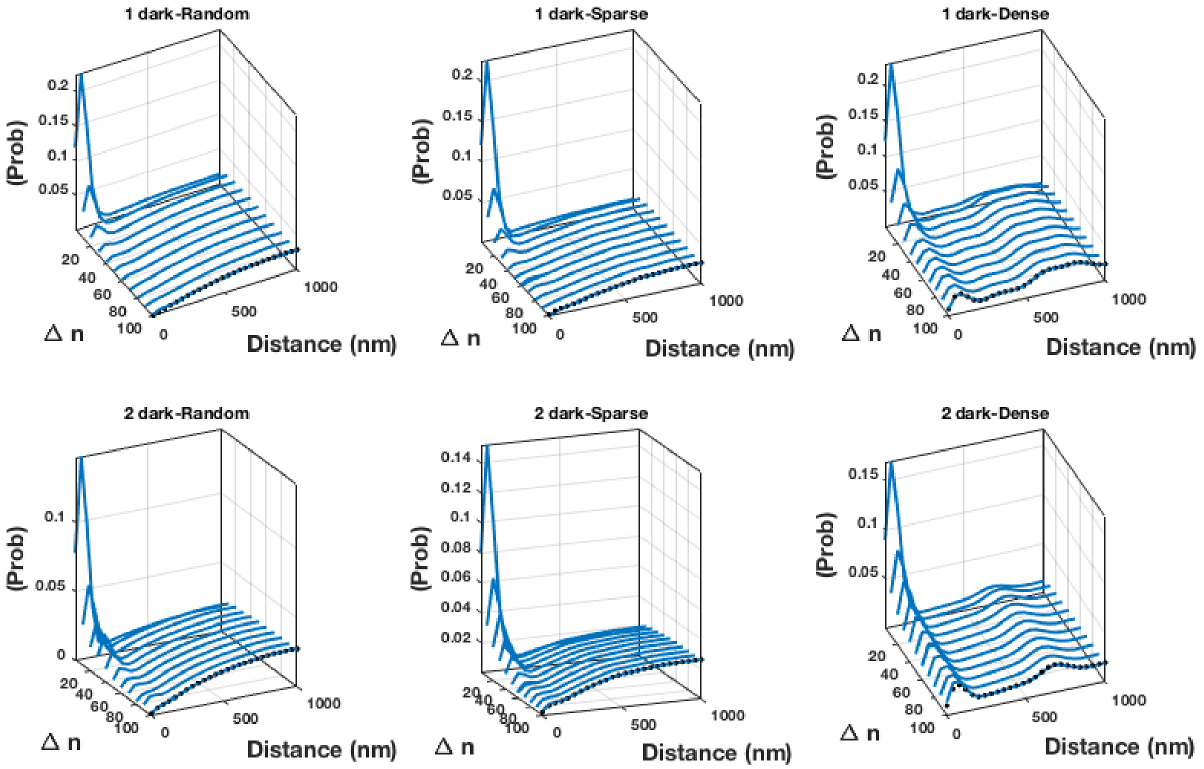

Figure S3: The pairwise distance distributions for both photo-kinetic models shown in Fig.S2 and 6 molecular assemblies. Note here that the axis is no longer log scale as in the main text and the true pairwise distance distribution is shown as black dots.

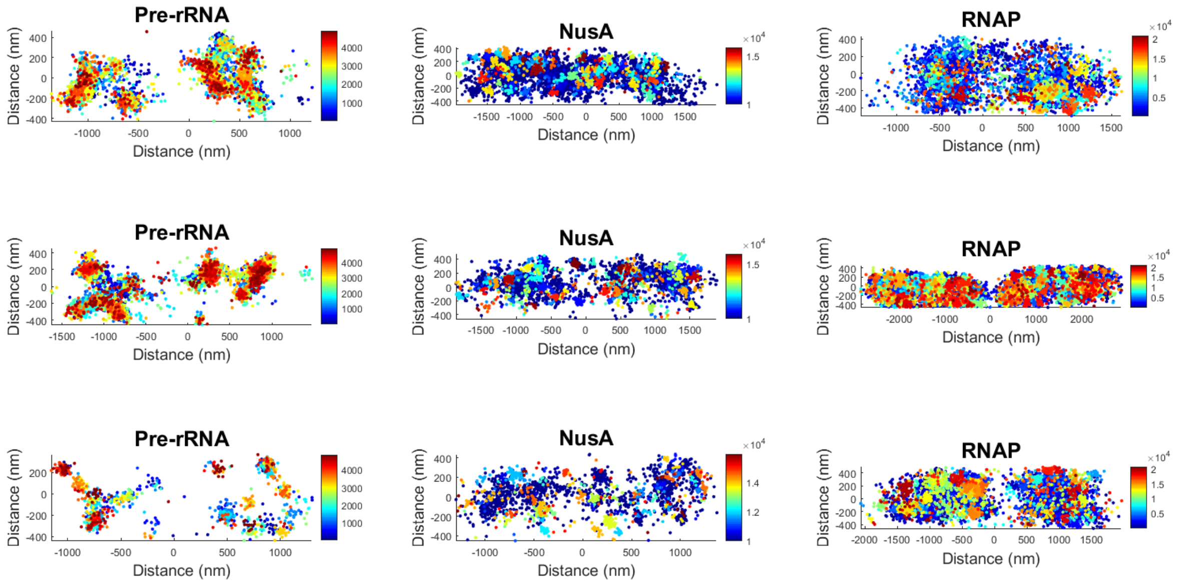

Figure S4: Example scatter plots of the experimental data used to verify that the pairwise distance distributions reached a steady state distribution. We show 3 cells for each molecular assembly, with the localizations colored with the frame of the localization.

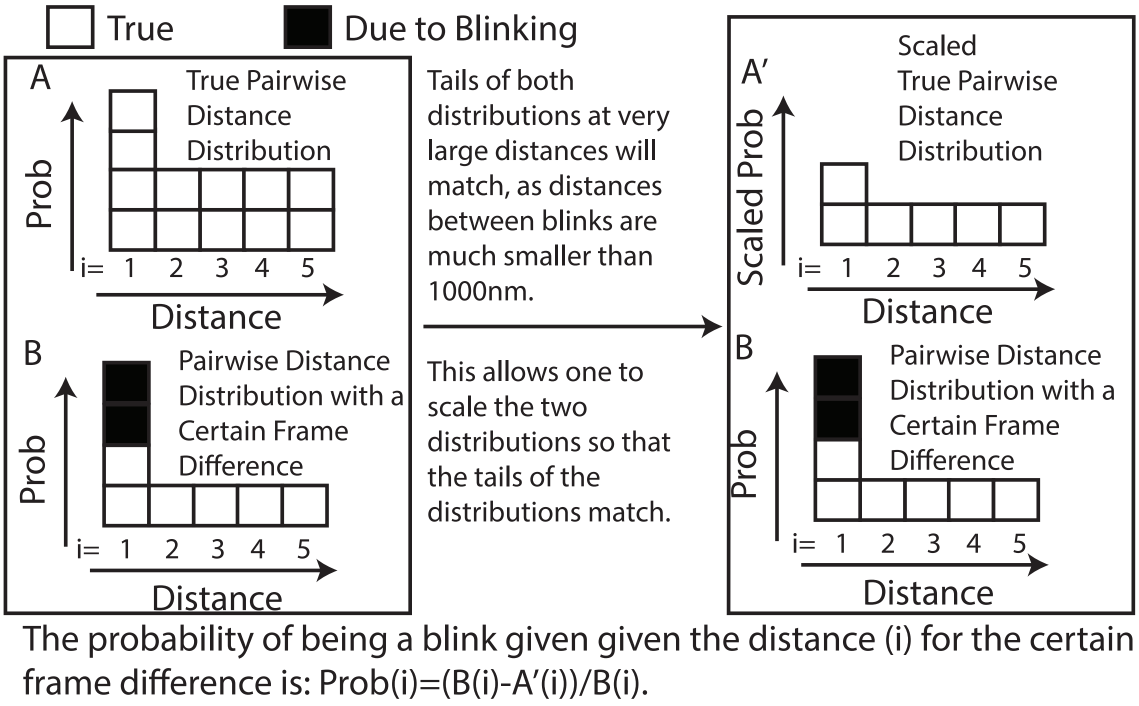

Figure S5: An illustration showing how to calculate $\mathbf{M}$ using the pairwise distance distributions. The blocks represent the distributions and $i$ is the distance bin.

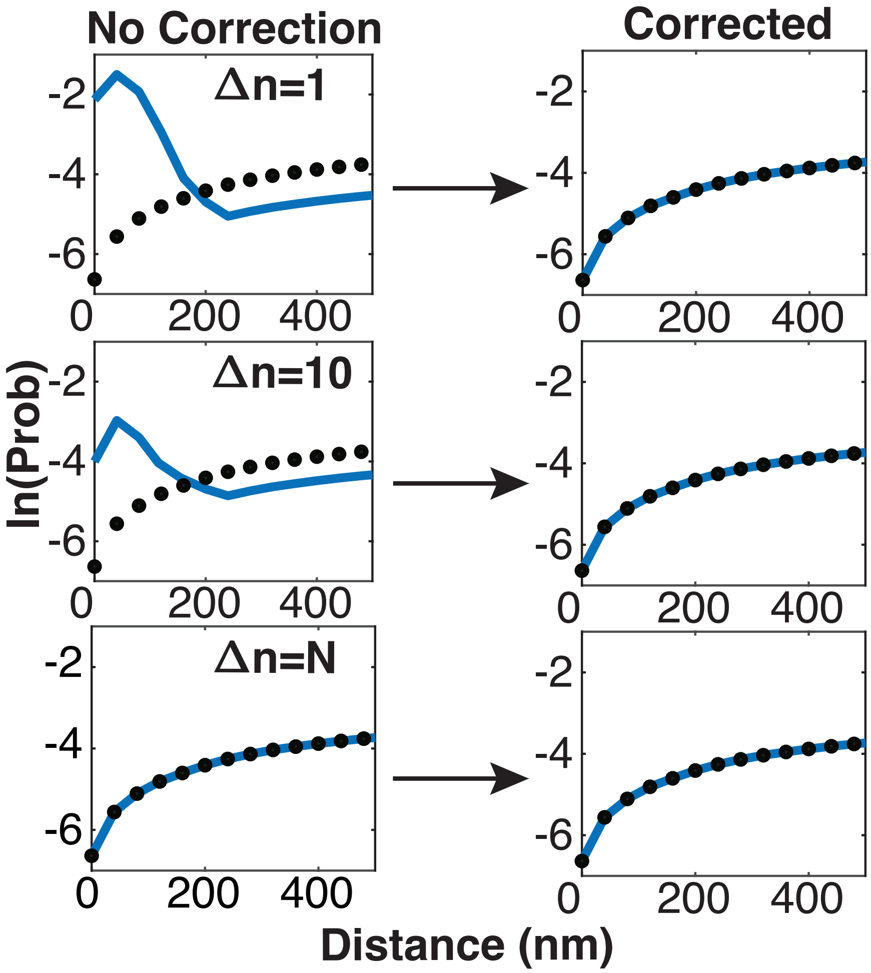

Figure S6: An illustration of the pairwise distance distributions at a certain frame difference, $\Delta n$, before and after being corrected with DDC. When the likelihood is maximized all of the pairwise distance distributions will match the true pairwise distance distribution. [The true pairwise distance distribution is shown as black dots.]

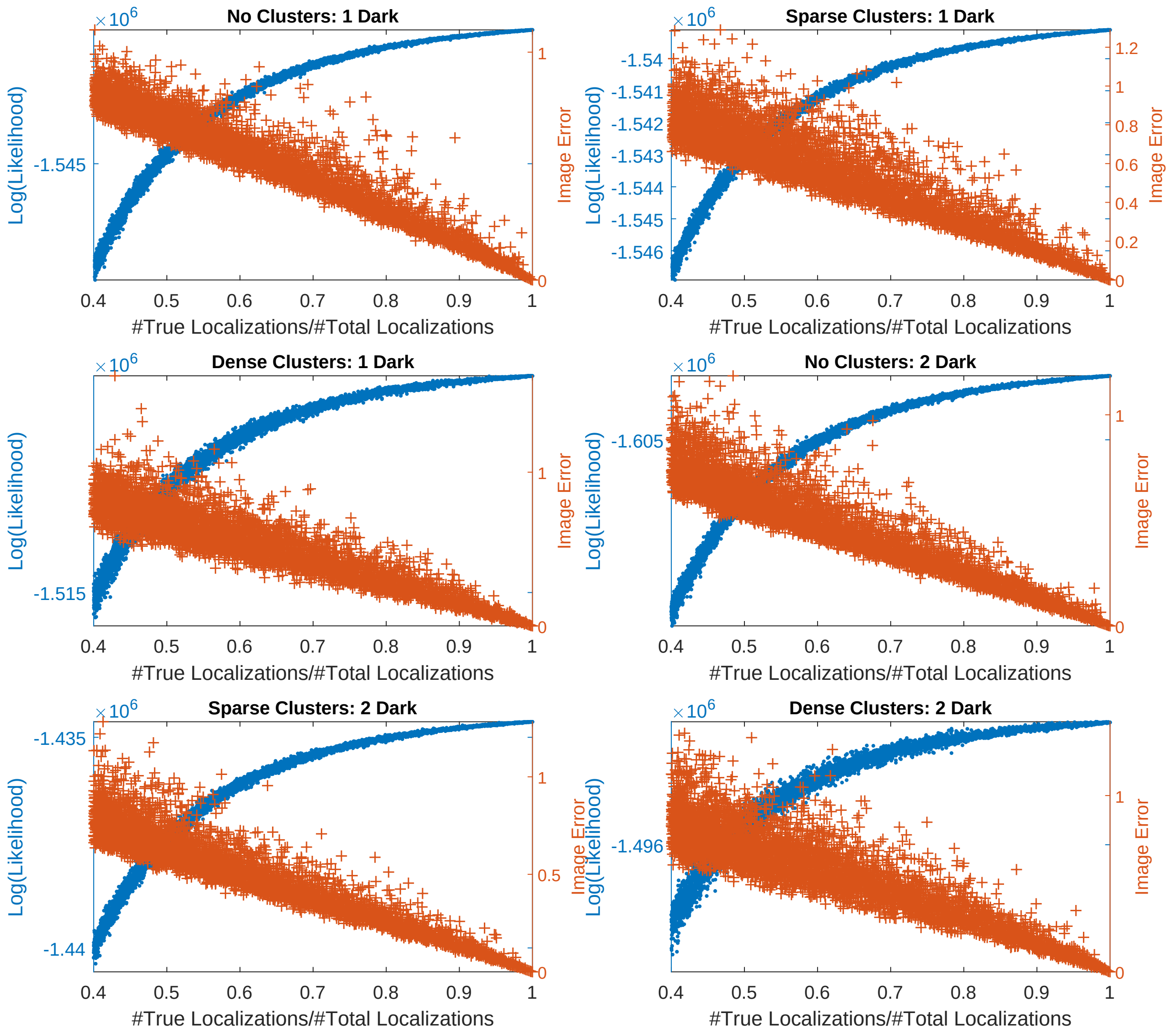

Figure S7: Maximization of Likelihood Results in Correct Conformation of Localizations: For 6 systems investigated within this work, we randomly varied the percentage of true localizations and calculated the $log(Lik)$ and the image error for each conformation (See Text).

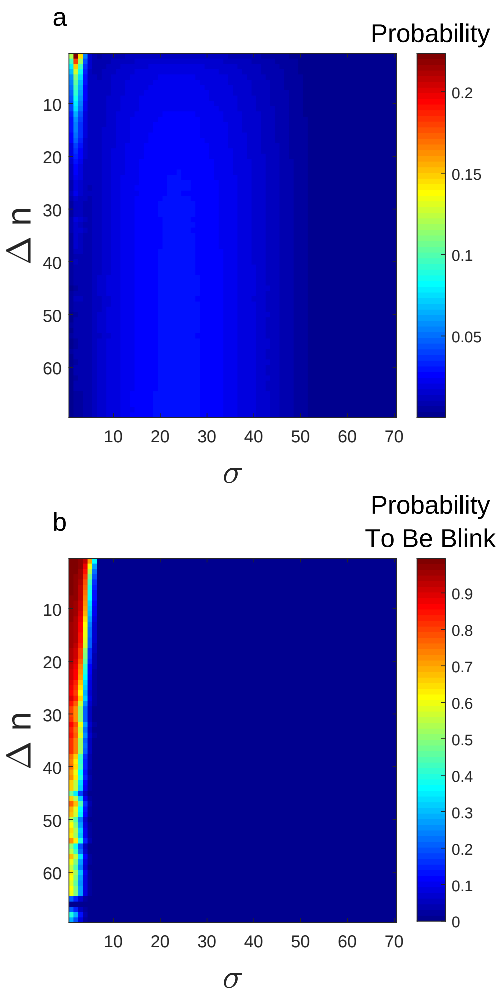

Figure S8: a. The probability distribution to observe a distance for a given $\Delta n$ , in units of resolution $\sigma$, between two localizations when at least one of them is a repeat, $P_{R1}(\Delta r|\Delta n)$. This specific distribution is for the 1 dark state no clusters system. (See Supporting Material text for details as to how these distributions are used to calculate Likelihood) b. The probability that a localization is the repeat of a given localization given the frame and distance between the localizations. These probabilities are calculated using the calculation shown in the prior figure.

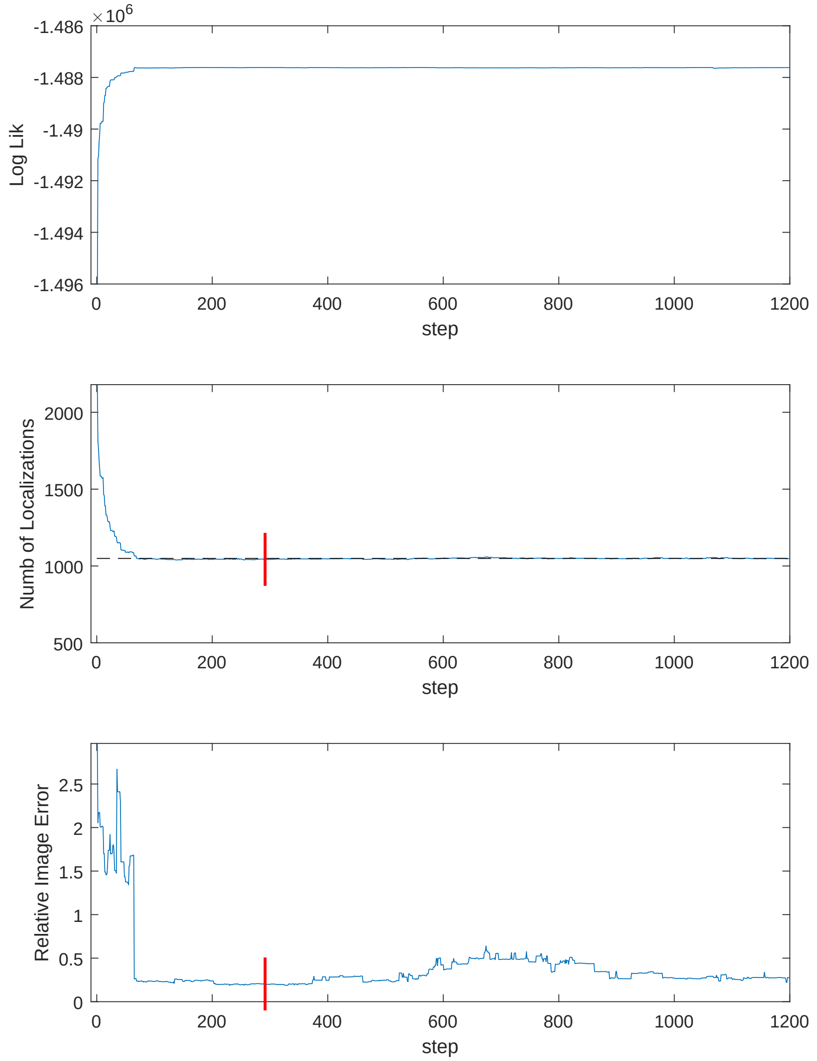

Figure S9: An example of the MCMC phase space search for the 2 dark state Small clusters system. For the number of localizations subplot a dashed black line shows the true number of localizations. For the bottom two subplots we show red lines indicating where the Likelihood was maximized. [Note: here we chose a random starting position for $\kappa(density)$ to illustrate the burn in phase of the MCMC, when $\kappa(density)$ starts at zero the burn in phase is not so extreme.]

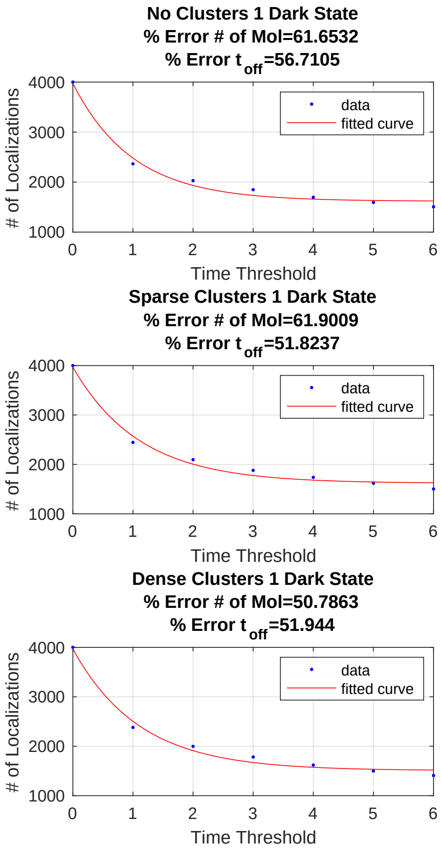

Figure S10: Resulting Error in Using Methodology of Annibale et al. (1): Here we only show the results for the 1 dark state systems with the fits to the semi-empirical formula (See Text). In the titles of each subplot we show the percent error in determining the number of true localizations and the average dark time.

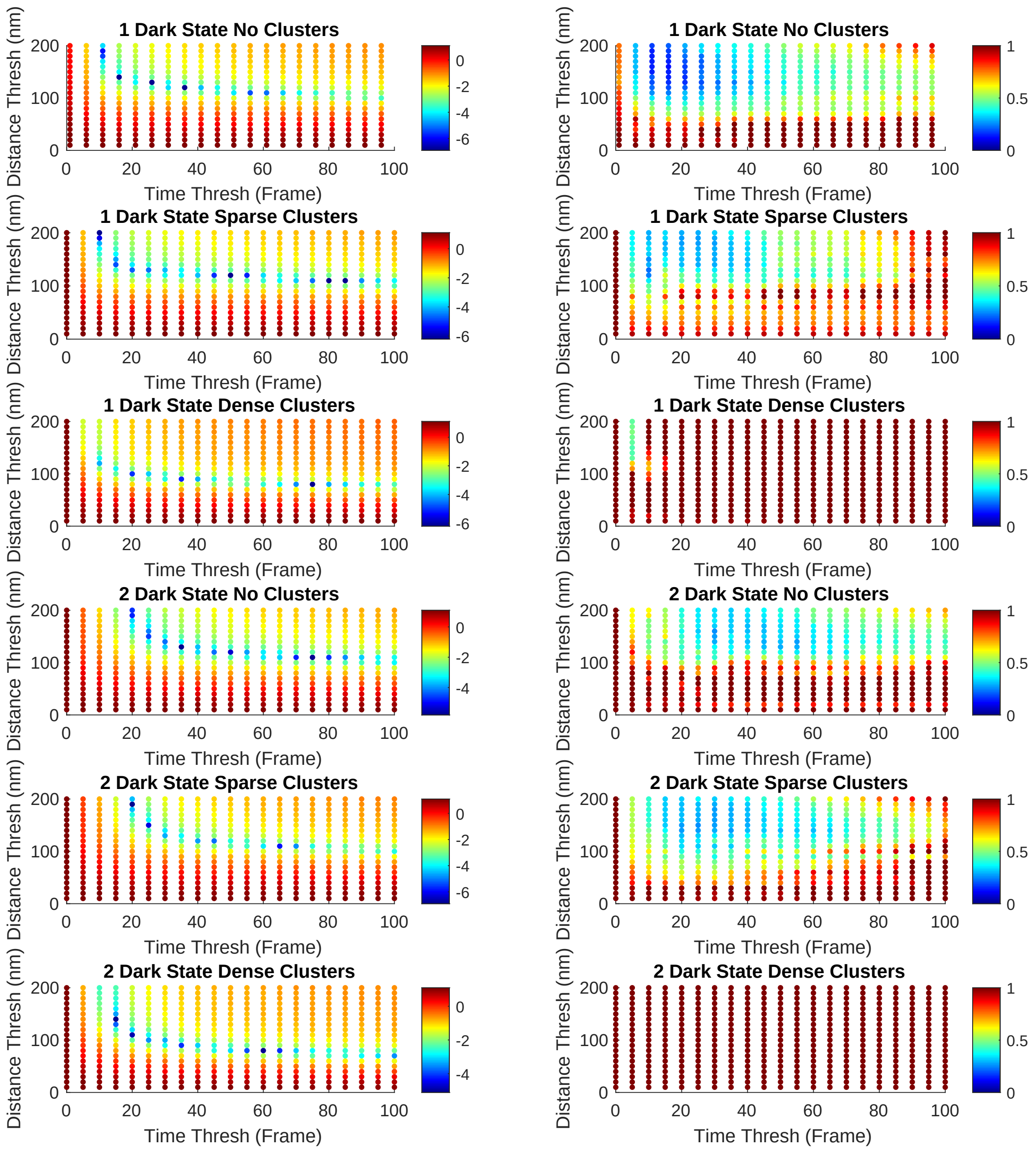

Figure S11: Determining the Thresholds for the Coltharp et al. Approach: In the first column we show the difference from the true number of localizations for the various time thresholds and distance thresholds, log scale ($ln[abs(\#loc - \#loc_{true})/\#loc_{true}]$. In the second column we plot the Image Error for each pair of threshold values for six systems.

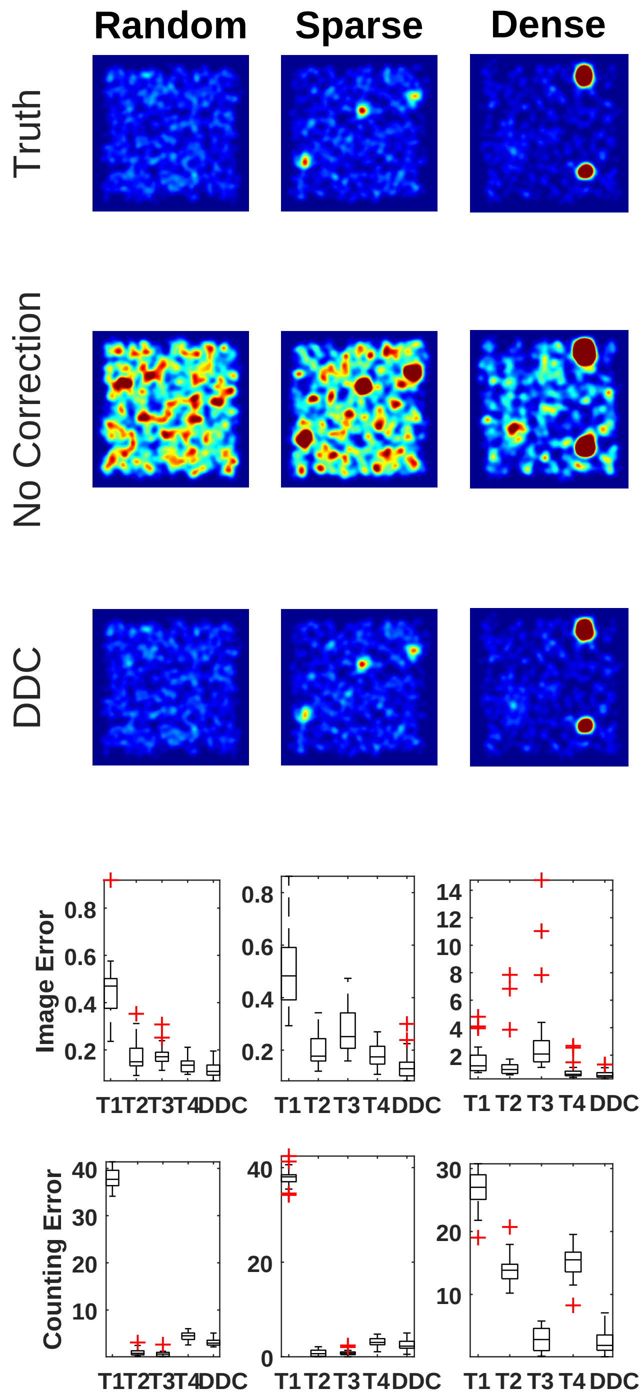

Figure S12: A comparison of the various thresholding methodologies with DDC and no blinking correction for the 1 dark state fluorophore. The first three rows show the images set to the same contrast for each labeled method. The last two rows show the results for the Image Error and the percent error in the number of fluorophores for each of the three systems for the one dark state fluorophore.

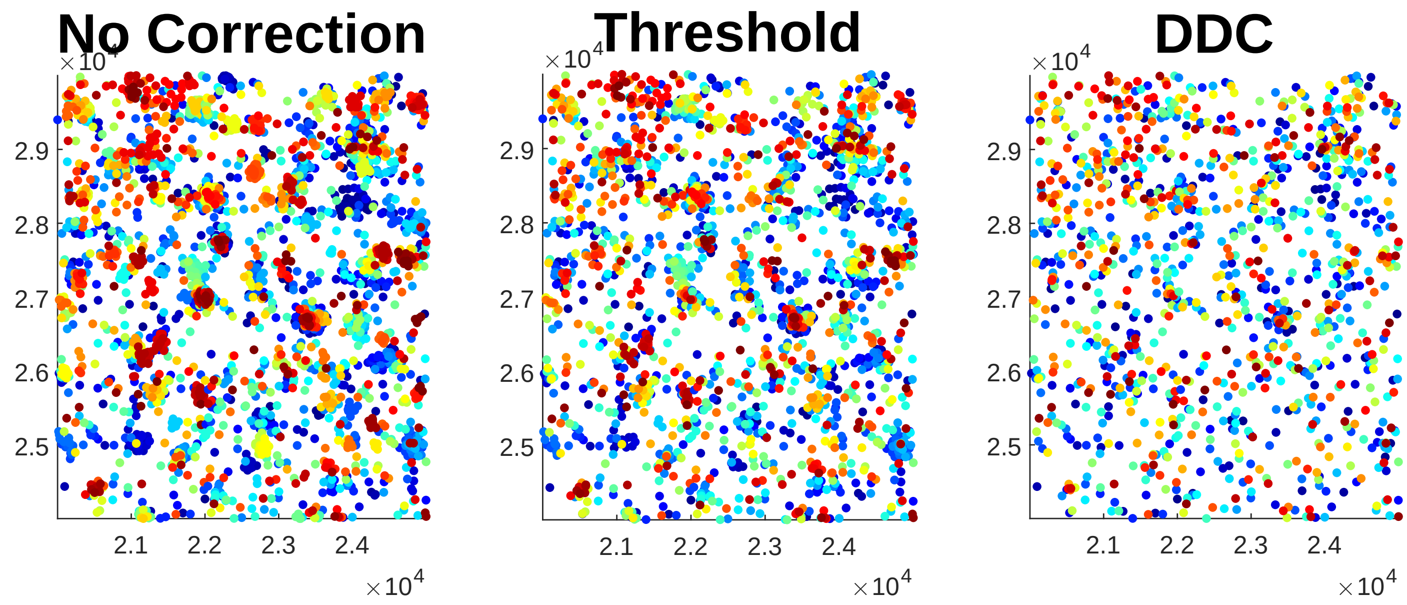

Figure S13: Scatter plots for a section of a cell with the localizations from AKAP79 with the color indicating the frame of the localization (Blue is early and Red is late). Here we show three different methodologies with the same thresholds used previously (19).

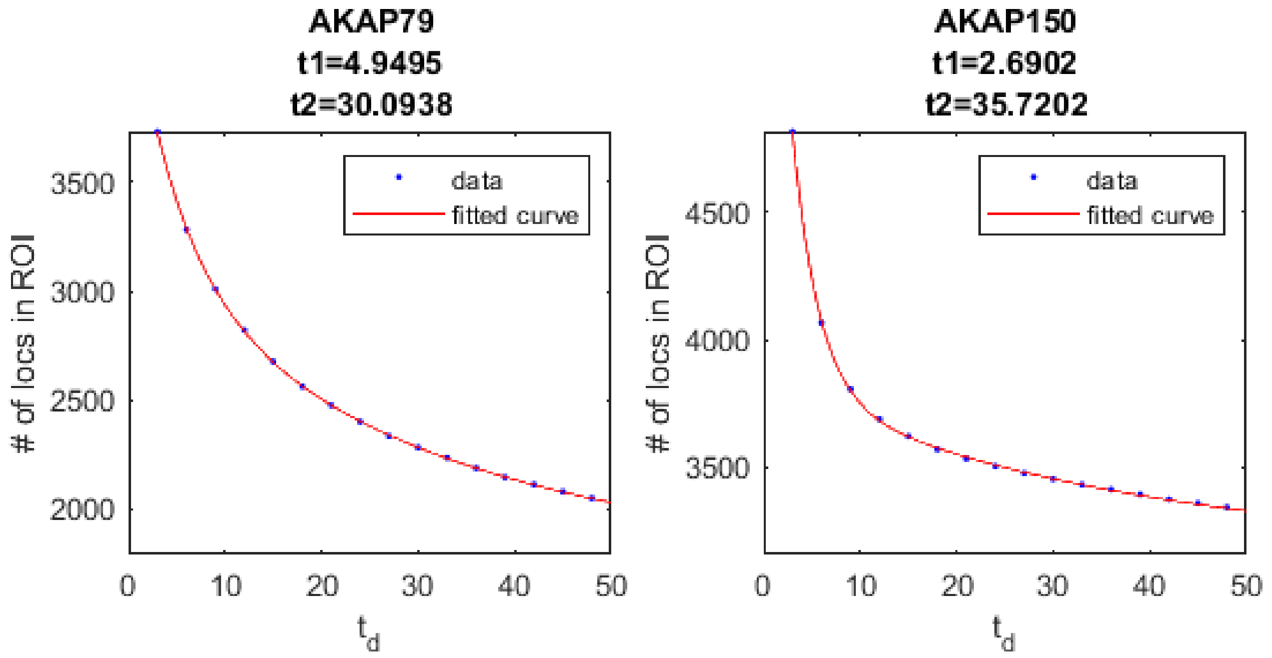

Figure S14: Here we show the results for determining the proper thresholds utilizing the methodology of T1 for AKAP79/AKAP150. The data was fitted to the double exponential used previously. Here the proper threshold is equal to two times the larger average dark time, either t1 or t2.

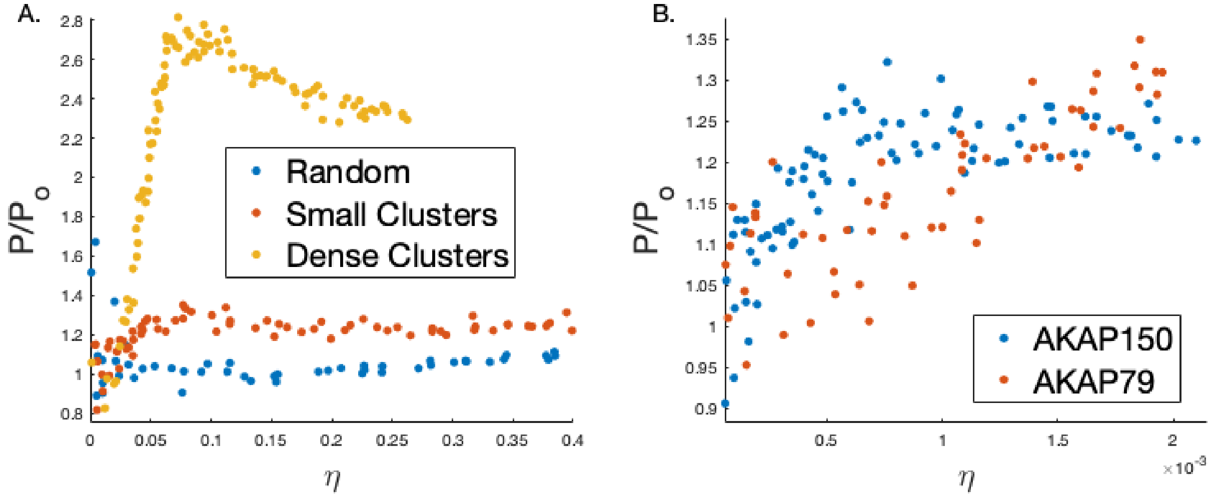

Figure S15: A. The results of computationally varying the label density on some of the simulation systems. B. The results of computationally varying the label density on AKAP79 and AKA150. (Values greater than 1 indicate significant clustering.)

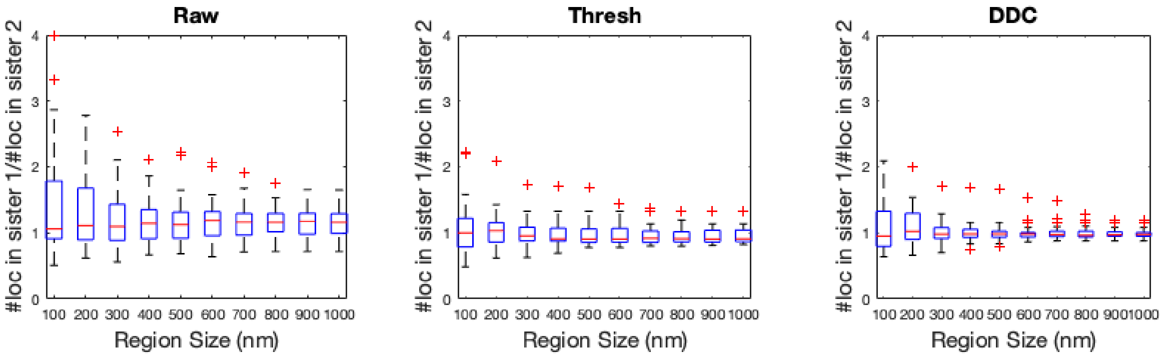

Figure S16: The ratio of the number of localizations between sister chromatids for each of the three methodologies using different sized segments along the fibers (Supporting Material, expected value is 1).

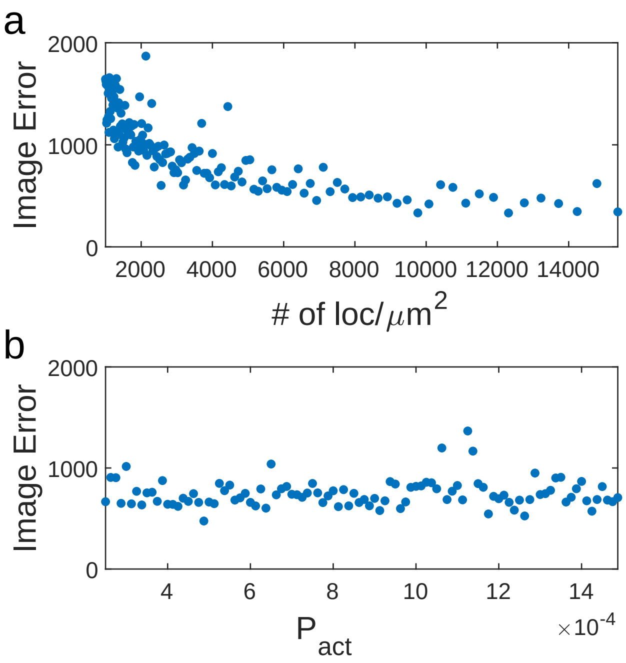

Figure S17: Here we show the raw Image Error (Not Normalized) for the uncorrected SMLM images for varying the density of the localizations and the activation energy.
